# A schizophrenia-associated radial glia cell state perturbs early human brain development

**DOI:** 10.1101/2025.10.24.684395

**Authors:** Susmita Malwade, Samudyata, Jessica Gracias Lekander, Angelika Langeder, Katharina Weinberger, Marja Koskuvi, Ana Osório Oliveira, Šárka Lehtonen, Luis Enrique Arroyo-García, Aiden Corvin, Sarah E. Bergen, I Ojansuu, O Vaurio, Jari Koistinaho, Jari Tiihonen, Carl M. Sellgren

## Abstract

Multidisciplinary evidence support a neurodevelopmental origin for schizophrenia, yet the mechanistic translation of known risk factors remains poorly understood. In this study, we leverage multi-lineage, forebrain-patterned organoids derived from monozygotic twins discordant for schizophrenia, integrating single-cell transcriptomic and epigenomic profiling to uncover disease-associated gene expression and chromatin accessibility topics. By constructing fate probability maps, we identify an accelerated developmental trajectory emerging from a distinct, schizophrenia-associated radial glia cell state, and reveal unique interactions among schizophrenia risk genes through unbiased multimodal analyses. Further, we confirm that schizophrenia-enriched states persist in differentiated lineages and disrupt synaptic programs, accompanied by molecular and cellular phenotypes mimicking observed disease pathology. Collectively, our findings delineate an early disruption in forebrain development, providing novel mechanistic insights into the origins of schizophrenia risk.

## INTRODUCTION

Schizophrenia (SCZ) is a severe psychiatric disorder typically diagnosed in late adolescence, when more diffuse prodromal symptoms converge into a recognizable spectrum of psychotic features^1^. However, recent studies assessing children at familial high risk for SCZ have also identified widespread neurocognitive impairments as early as age two^2,3^, underscoring the neurodevelopmental origins of the disorder. SCZ affects an estimated 24 million people worldwide^4^, and is primarily driven by genetic risk factors, with heritability estimates ranging from 60–80%^5,6^. The genetic architecture of SCZ is highly polygenic^7^, and genome-wide association studies have identified over 300 loci associated with the disorder ^8^. Despite these findings, translating genetic risk into mechanistic insight has remained challenging. A major limitation is the scarcity of disease-relevant brain tissue from early developmental stages— periods strongly implicated in SCZ by both temporal gene expression patterns and the enrichment of high-risk variants^7–12^.

Human brain organoid models offer a promising alternative by recapitulating the dynamic cytoarchitecture of the developing human brain while preserving patient-specific genomic context^13–15^. However, capturing the polygenicity of SCZ at the required scale remains difficult, at least if relying on randomly sampled cases. Alternatively, engineering isogenic lines with introduced risk variants raises concerns about whether these constructs faithfully model the complexity of the disorder.

To address these limitations, we generated multi-lineage, forebrain-patterned organoids from monozygotic twin pairs discordant for SCZ, where one twin developed the disorder while the co-twin remained unaffected beyond the age of typical onset. This unique design theoretically enabled the interrogation of *de novo* variants sufficient to drive disease phenotypes within an otherwise shared genomic background.

We then applied single nucleus multiome (RNA + ATAC) sequencing and probabilistic topic modeling to define molecular programs associated with SCZ. Fate probability mapping revealed an accelerated developmental trajectory originating from a distinct and SCZ-associated radial glial state, marked by unique regulatory interactions among SCZ risk genes. This pathological state persisted across differentiated lineages and included synaptic programs with molecular and cellular consequences consistent with hallmark features of the disorder, including synaptic loss^16,17^.

## RESULTS

### Multiomic characterization of the forebrain patterned organoids

Induced pluripotent stem cells (iPSCs) derived from three monozygotic twin pairs discordant for schizophrenia (MZ_d_) were used to generate forebrain-patterned brain organoids directed towards anterior neuroectodermal fate^18^ (Fig. 1a, Supplementary Table 1). To also capture the impact of microglia on developing neuronal circuits^19–21^, we incorporated primitive macrophage progenitors (PMPs) generated in a Myb-independent manner^22^, recapitulating the early developmental infiltration of microglia^23^. Their maturation within the organoid environment was supported by supplementation with IL-34 and GM-CSF (Fig. 1b, Methods, Extended Data Fig. 1a–b).

**Figure 1.**
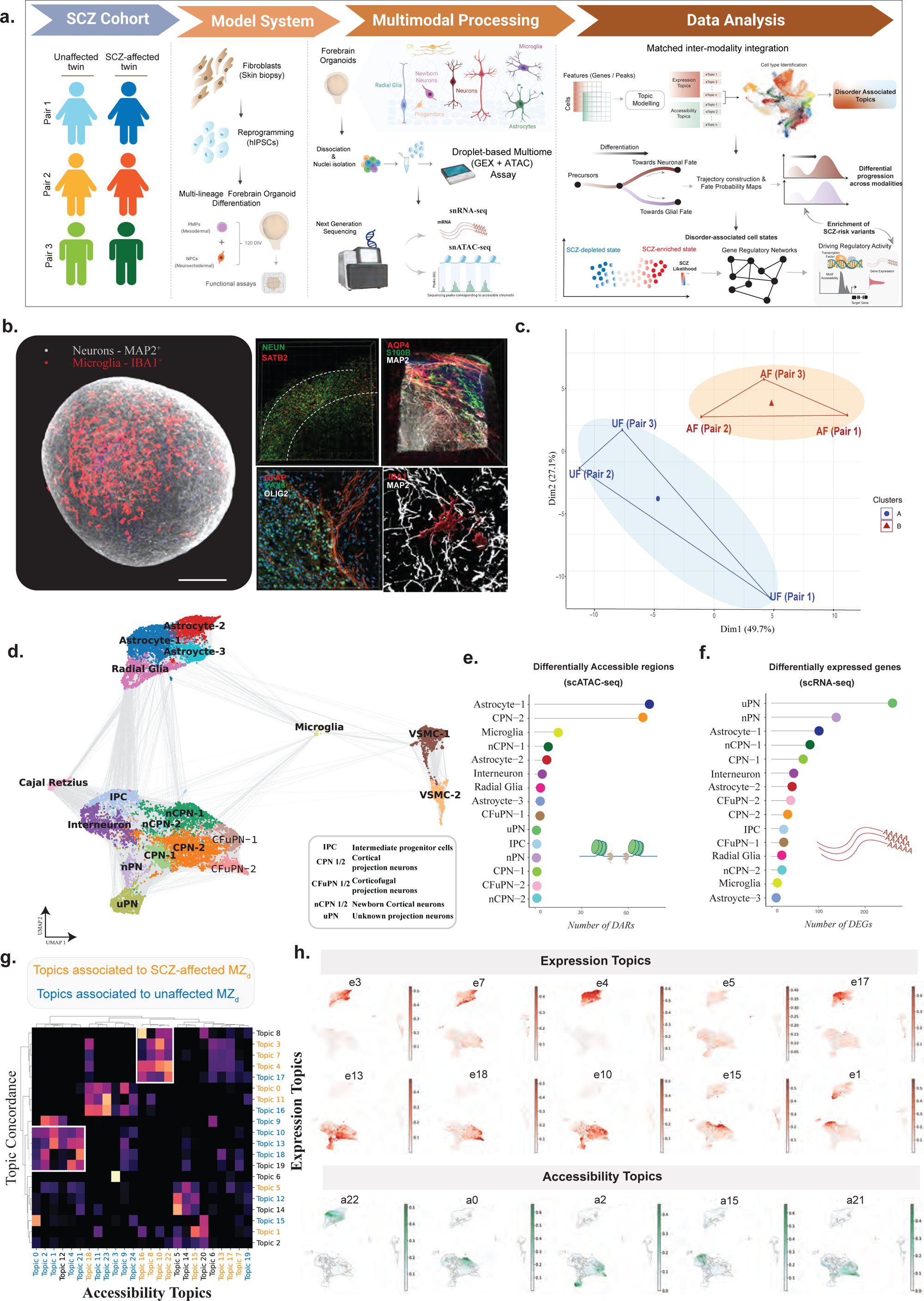
**(A)** Schematic representation of the experimental and computational workflow of this study. **(B)** Confocal microscopy images of optically cleared whole organoid showing the presence and distribution of neurons (MAP2) and microglia (IBA1) within the organoid (left). Immunohistochemical staining of neurodevelopmental markers at DIV 120 for neurons (NEUN; SATB2; MAP2), astrocytic lineage (S100B, AQP4, GFAP), neuronal precursors (PAX6), microglia (IBA1), and oligodendrocyte-lineage (OLIG2). White dashed lines indicate formation of the cortical plate at the periphery of the organoid. Scale bar represents 500μm. (Additional markers are provided in Extended Data Fig. 1c). **(C)** Principal component analysis (PCA) plot of bulk proteomic abundance data from patient-derived organoids of MZ_d_ twins (n=6 subjects). Each dot represents a subject colored by condition (unaffected or SCZ-affected twin). **(D)** UMAP embedding of joint RNA and ATAC modality KNN graph displaying 18 unique cellular clusters. **(E)** Cell type-specific number of differentially expressed genes (DEG) from the RNA modality per cluster (left), and **(F)** differentially accessible regions (DAR) from the ATAC modality per cluster (x-axis) represented as a bar graph. **(G)** Heatmap displaying concordance between expression topics (y-axis) and accessibility topics (x-axis) across the computed cell-topic matrix. Highly concordant topics indicate plausible stable cell states (white box). Topics associated to SCZ-affected MZ_d_ are colored in orange and unaffected MZ_d_ in blue. **(H)** UMAP plot displaying the distribution of selected topics with expression topics (red) and accessibility topics (green) across cells. Cells are colored by topic activation values.

After 120 days in vitro (DIV), we validated the organoid model by confirming the presence of neural progenitors, neurons, astrocytes, and microglia, along with both pre- and post-synaptic structures, using optical tissue clearing followed by immunohistochemistry (Fig. 1b, Extended Data Fig. 1c). Electrophysiological activity was verified prior to downstream processing using whole-cell patch-clamp recordings (Extended Data Fig. 1d).

As a proof of concept that the organoid model could resolve SCZ-relevant molecular signatures despite the near-identical genetic background within each discordant monozygotic twin pair, we performed mass spectrometry-based proteomic profiling on 120-day-old organoids (DIV 120; n=3 subjects per diagnosis). This confirmed that proteomic profiles clustered by diagnostic status rather than by twin pair, supporting disease-associated molecular divergence (Fig. 1c).

We then pooled six organoids per donor (DIV 120; *n* =6 subjects) and performed single-nucleus multiome (RNA + ATAC) sequencing to simultaneously capture transcriptomic and chromatin accessibility profiles of each cell (Fig. 1a). Rigorous quality control yielded 21,745 high-quality nuclei with paired gene expression and accessibility peak counts.

We then applied probabilistic topic modeling integrated with variational autoencoders, a framework recently developed for single-cell multiomic analysis¹⁷ (Fig. 1d, Extended Data Fig. 2a–i). This approach, optimized using Bayesian hyperparameter tuning, identifies transcriptional and regulatory topics—each corresponding to groups of co-expressed genes or co-accessible cis-regulatory elements—and infers cell states as mixtures of these topics (Extended Data Fig. 3a–b). All identified topics contributed to a joint k-nearest neighbor (KNN) graph embedded in a low-dimensional space, revealing 18 transcriptionally and epigenetically distinct cellular clusters across donors (Fig. 1d).

Clusters were evaluated for donor and technical batch effects (Extended Data Fig. 2e– f), and annotated based on projection onto reference fetal forebrain atlases and canonical marker signatures. Cell types included radial glia, intermediate progenitors, multiple neuronal subtypes (newborn neurons, corticofugal neurons [CFuPN], cortical projection neurons [CPN], interneurons, and unspecified projection neurons), astroglial subtypes, microglia, and mesenchymal progenitors (Methods; Fig. 1d; Extended Data Fig. 4a–d; Supplementary Table 2). Overall, organoids at DIV 120 exhibited transcriptomic profiles broadly resembling human fetal forebrain from the second trimester (Extended Data Fig. 4c).

### Multimodal topics define neurodevelopmental cellular states in schizophrenia

We then analyzed each modality independently, performing paired comparisons across twin pairs to identify cell type–specific differentially expressed genes (DEGs) and differentially accessible regions (DARs) (Fig. 1e–f; Supplementary Table 3). Astrocyte and neuronal subtypes exhibited the most pronounced epigenetic alterations, while differentiated neurons showed marked transcriptomic changes, particularly in genes involved in cell adhesion and axon guidance (Fig. 1e–f).

To assess the convergence of modalities in relation to the disease, we evaluated associations between expression and accessibility topics at the single-cell level using mixed-effect regression models that accounted for twin pairing. Topics significantly associated with disease status exhibited greater cross-modality concordance, reflecting more distinct and transcriptionally coherent cell states (Fig. 1g; Supplementary Table 4). Among the SCZ-associated topics (e3, e7, e4, e17, a22, a10, a8, a16; *P_adj_* < 0.05), we observed predominant enrichment in radial glia and astrocytic clusters within the joint latent space. These topics captured key biological processes such as ‘immune activation’ (e3), ‘cell cycle progression’ (e7), ‘forebrain cell migration’ and ‘dendritic morphogenesis’ (a22), as well as ‘phagocytosis’ (e5) in microglia (Fig. 1g–h; Supplementary Table 5). In contrast, topics enriched in unaffected co-twins were distributed primarily across neuronal clusters. These included expression topics e13 and e18, associated with ‘neuron projection development’ and ‘calcium ion homeostasis’, respectively, and accessibility topics a0 and a21, linked to ‘cell fate commitment’ (*P_adj_* < 0.05; Fig. 1g–h; Supplementary Table 5).

### Premature lineage commitment in schizophrenia

To further investigate the mechanistic underpinnings of the identified topics, we reconstructed a joint KNN graph of telencephalic lineage cells and inferred a pseudotime-ordered cell-state tree represented as a dendrogram (Fig. 2a). As expected for the developmental window examined, the trajectory recapitulated canonical lineage bifurcations: radial glia and intermediate progenitors diverging into neuronal and glial lineages (Fig. 2b). When comparing discordant twins, we observed significant disruptions in both lineage differentiation (*P* = 4.8 × 10⁻⁵) and developmental progression (*P* < 1 × 10⁻⁶) in SCZ twins, characterized by an earlier acquisition of gliogenic competence (Fig. 2d–e).

**Figure 2.**
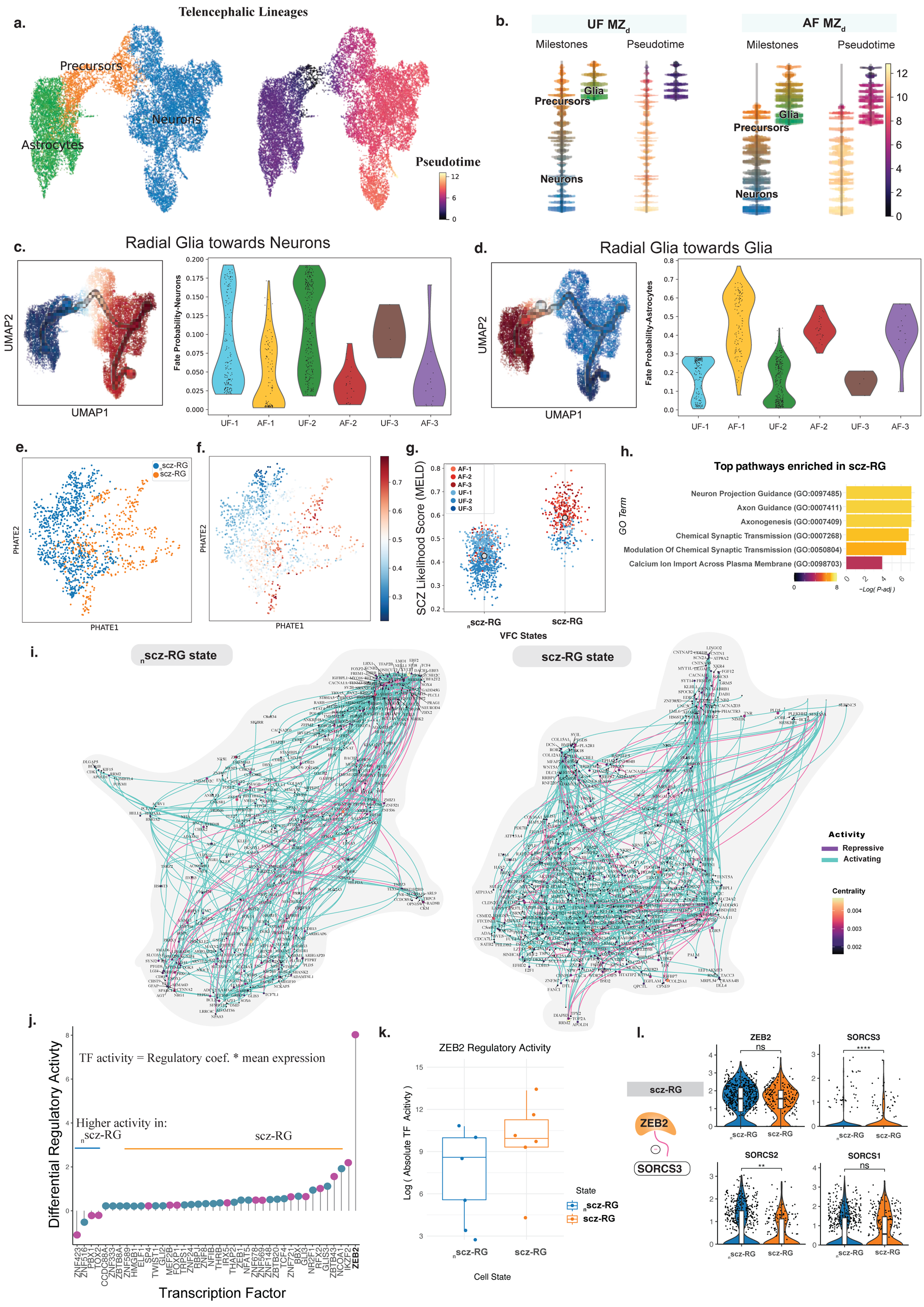
**(A)** UMAP embedding of cells belonging to telencephalic lineages (all radial glia, intermediate progenitors, neuronal, and astrocytic clusters) upon recomputing the joint KNN graph. Cell are coloured by the lineage (left; radial glia and intermediate progenitors labelled as precursors) and by computed pseudotime values (right) with black dots representing terminal states inferred by CellRank. **(B)** Dendrogram of cells of unaffected twins (top) and SCZ-affected twins (bottom) representing the bifurcation analysis (scFates). Cells are colored by assigned milestones (left) based on pseudotime (right) individually. Fate probabilities of radial glia cells (y-axis) across all MZ_d_ (x-axis) towards the **(C)** neuronal terminal states and **(D)** astrocyte terminal states visualized on UMAPs (left) and violin distributions (right). [Neuronal fate probabilities per twin pair: P_pair 1_ = 1.02e-09, P_pair 2_ = 3.67e-11, P_pair 3_ = 0.0092; Mann-Whitney U test]. **(E)** PHATE embedding of radial glia (RG) cluster with cells colored by the identified vertex frequency subclusters and **(F)** SCZ-likelihood scores. **(G)** Jitter plot displays cells belonging to each MZ_d_ subject across both RG cell states (x-axis) and their SCZ-likelihood scores (y-axis). RG cell state enriched in SCZ-affected MZ_d_ is labelled as scz-RG and unaffected MZ_d_ is labelled as _n_scz-RG. Legend indicates the MZ_d_ twin subject. **(H)** Top enriched pathways (GO:BP terms) in the scz-RG state compared to _n_scz-RG state (P-adjusted<0.05). **(I)** Gene regulatory networks (GRN; RNA+ATAC) of the SCZ-enriched RG cell state (right) and unaffected MZ_d_ -enriched RG cell state (left) computed by fitting generalized linear regression models for transcription factors (TFs), TF-binding motif accessibility, and expression of target genes, implemented in Pando. The TF networks are visualized on a UMAP embedding based on coexpression and inferred interaction strength between TFs. Color and size of the dot represent PageRank centrality values of each significant TF (node; P-adjusted<0.05). Edges are colored by activity of the TF inferred by the sign of the regulatory coefficient (repressive-pink, activating-cyan). All significant TF-accessible peak-target gene trios of the GRNs are provided in Supplementary Table 6. **(J)** Top TFs with differential activity between the two RG cell states (scz-RG versus _n_scz-RG). Activity for each TF is calculated as mean regulatory coefficient*mean expression of the TF. Differences in activities of each TF is shown on the y-axis with TFs on the x-axis. **(K)** Top transcription factors and their inferred target genes with varying regulatory activity in GRNs of scz-RG (right) versus _n_scz-RG (left) cell states. Magenta indivates repressive activity and cyan indicates activation.

### A disease-associated radial glia subpopulation

Given that radial glia serve as the primary predecessors to neurogenesis and gliogenesis, we next examined whether this population exhibited differential fate selection in SCZ. We computed fate probability maps and traced terminal differentiation states originating from radial glia (Extended Data Fig. 5a). Unsupervised inference of genes driving each fate yielded known regulators of lineage specification, validating the inferred maps (Fig. 2b; Extended Data Fig. 5a–b). Consistent with the accelerated developmental trajectories observed in SCZ, radial glia from twins with SCZ displayed significantly higher fate probabilities toward glial lineages compared to their unaffected co-twins (Fig. 2c). Despite this skewed commitment, the overall composition of terminally differentiated cell types remained relatively stable (Extended Data Fig. 5c).

To further dissect this bias, we isolated radial glia and estimated per-cell SCZ-likelihood estimates to define SCZ-enriched (scz-RG) and depleted (_n_scz-RG) subpopulations (Methods, Fig. 2d–f; Extended Data Fig. 5d). We then inferred gene regulatory networks (GRNs) across the two radial glial states using linear modeling of transcription factor (TF) binding motifs and their target gene expression^24^ (regulatory coefficient * mean expression) (Fig. 2g–i; Supplementary Table 6).

Differential TF activity inferred from GRNs revealed that *ZEB2*—a key regulator of early neocortical cell fate and gliogenesis^25^, and an established SCZ risk gene^7,26,27^—exhibited significantly elevated regulatory activity in the scz-RG subpopulation compared to the _n_scz-RG subpopulation (Fig. 2j; Supplementary Table 6). Notably, this increase in regulatory activity occurred without corresponding changes in *ZEB2* transcript levels (Fig. 2k). This pattern was consistently observed across all twin pairs, implicating altered *ZEB2*-mediated transcriptional control as a potential upstream driver of the gliogenic bias observed in SCZ.

Further analysis of *ZEB2* target gene networks revealed functional divergence between the SCZ-enriched and SCZ-depleted radial glia states. In the scz-RG subpopulation, *ZEB2* was predicted to repress *SORCS3*, a sortilin-related synaptic receptor and established SCZ-risk gene^9^, while in the _n_scz-RG subpopulation, *ZEB2* was inferred to function as an activator of *SEMA3A*, another SCZ risk gene that encodes an axon guiding cue^28–30^ (Supplementary Table 6).

To also assess potential downstream consequences of altered RG states, we performed unsupervised intercellular communication inference from RG to neurons and astrocytes. This analysis identified disrupted NRG3–ERBB4 signaling in SCZ-affected twins (*P* < 0.05; Extended Data Fig. 5e)— a pathway previously implicated in SCZ^10,31,32^ and known to regulate both cortical plate invasion²² and astrocyte-mediated synaptogenesis²³. Combined, these findings suggest that early transcriptional dysregulation in radial glia may propagate to later deficits in neuronal migration and glial support mechanisms critical for cortical development.

### Downregulation of the scz-RG associated SORCS proteins across modalities and cohorts

Guided by the GRN analyses, we next examined whether inferred driver genes or their downstream targets exhibited consistent signatures across other molecular modalities. Focusing on *SORCS3*, the predicted *ZEB2* target in scz-RG, we also evaluated its closely related homologs, *SORCS1* and *SORCS2*^33^, members of the sortilin-related VPS10P domain-containing receptor family. We observed reduced mRNA expression of both *SORCS2* and *SORCS3* in the scz-RG cell population compared to _n_scz-RG cells (Fig. 2l). Although *SORCS3* protein levels were below detection in our proteomic dataset, *SORCS2* was significantly downregulated in organoids derived from affected twins relative to their unaffected co-twins (Extended Data Fig. 1e).

To assess the generalizability of our findings, we analyzed whole-genome sequencing (WGS) data from an independent cohort of nine monozygotic twin pairs discordant for SCZ. We identified 56 discordant copy number variation (CNV) regions—46 deletions and 10 duplications—predominantly located in intergenic regions and present in at least four discordant twin pairs (Extended Data Fig. 5f; Supplementary Table 14). Notably, one recurrent deletion mapped to intron 2 of *SORCS2,* appearing in four of the nine pairs. Complementary analysis of postmortem brain tissue revealed that *SORCS2* and *SORCS3* share transcriptional networks during brain development, significantly enriched for synaptic organization and signaling pathways (*P_adj_* < 0.05; Extended Data Fig. 5g).

Additionally, we identified several discordant regions associated with genes that are highly expressed during prenatal brain development and play key roles in neurogenesis. These included *MCPH1*, linked to premature neuronal differentiation^34^, *LINGO2*^35^, and *TENM1*, involved in neuronal connectivity^36^; and *RGS12*, regulating neuronal signalling^37^.

### Synaptic consequences along the neuronal trajectory

Given the identified SCZ-enriched radial glia state, we investigated the impact on gene expression as development progressed along the neuronal trajectory. To do this, we fitted negative binomial generalized additive models (GAMs) for each gene and TF motif across pseudotime.

Along this neuronal trajectory, we identified 164 significantly dysregulated genes (*P_adj_* < 0.05), with enrichment at the postsynapse (Fig. 3a–b), suggesting disrupted neurite projection and migration (e.g., *RIT2*, *CTNNA2*, *OLFM3*, *SEMA3D*, *FSTL4*, *PTPRO*, *LHX1*, *BCL11B*, *RELN*), cell adhesion and junction organization (*PCDH15*, *FREM1*, *FLRT2*), and synaptic signaling (*SYN3*, *GRIK2*, *GABRB2*, *CNTNAP4*, *CHRNA7*) (*P_adj_* < 0.05; Fig. 3c, Supplementary Table 7). Trajectory-associated topic modeling across twin pairs (diagnosis × pseudotime) revealed enriched biological themes including cell division (topic e8) and cell fate commitment (topics a11, a21) in precursor cells, as well as cytoplasmic translation (e14), synapse assembly (e19), and calcium ion transport (e18) in neurons (Fig. 3d–e; Supplementary Table 8).

**Figure 3.**
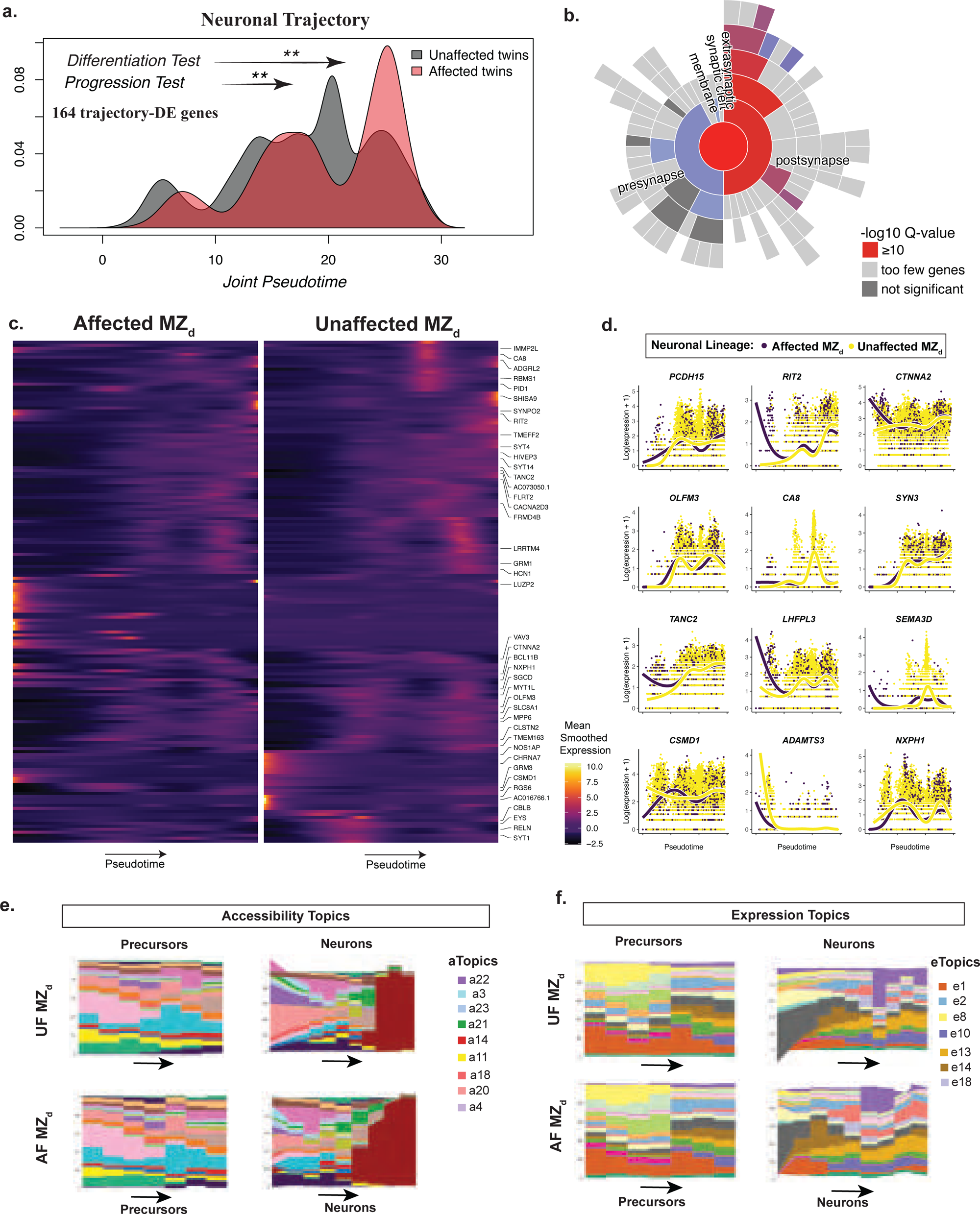
**(A)** Density distribution of cells belonging to each condition along the progression of neuronal lineage as indiciated by pseudotime (y-axis). **(B)** Sunburst plot displaying the enrichment of neuronal trajectory-associated differentially expressed genes in post-synapse gene sets obtained from SynGO database (one-sided Fisher exact test; FDR<0.05). **(C)** Heatmap showing mean smoother expression values of significant trajectory-associated differentially expressed genes (pseudotime*condition) in the neuronal lineage across pseudotime (y-axis) (P-adjusted < 0.05). Genes are clustered using heirarchical clustering. **(D)** Scatter plot of lof-transformed normalized expression (y-axis) of selected trajectory-associated DEGs across pseudotime (x-axis) in the neuronal lineage. Cells are coloured according to the condition as shown in the legend. Colored line represents a smoothing spline fit in the respective condition. **(E)** Streamgraphs of accessibility topics distributed along pseudotime (y-axis) in precursors and neuronal clusters comparing cells from unaffected twins (top) to SCZ-affected twins (bottom). Color indicates accessibility topic number. **(F)** Streamgraphs of expression topics distributed along pseudotime (y-axis) in precursors and neuronal clusters comparing cells from unaffected twins (top) to SCZ-affected twins (bottom). Color indicates expression topic number.

### Increased uptake of synaptic material in microglia

Among the topics significantly associated with SCZ, we also identified a set of phagocytic genes defining topic *e5* in microglia (*P_adj_* < 0.05; Fig. 4a–b, Supplementary Table 5). Notably, reduced synapse density is a well-documented feature early in the course of SCZ^38,39^, and prior studies using 2D cellular models derived from SCZ patients have shown excessive microglial uptake of synaptic structures, an effect driven by both synapse- and microglia-related factors^17^.

**Figure 4.**
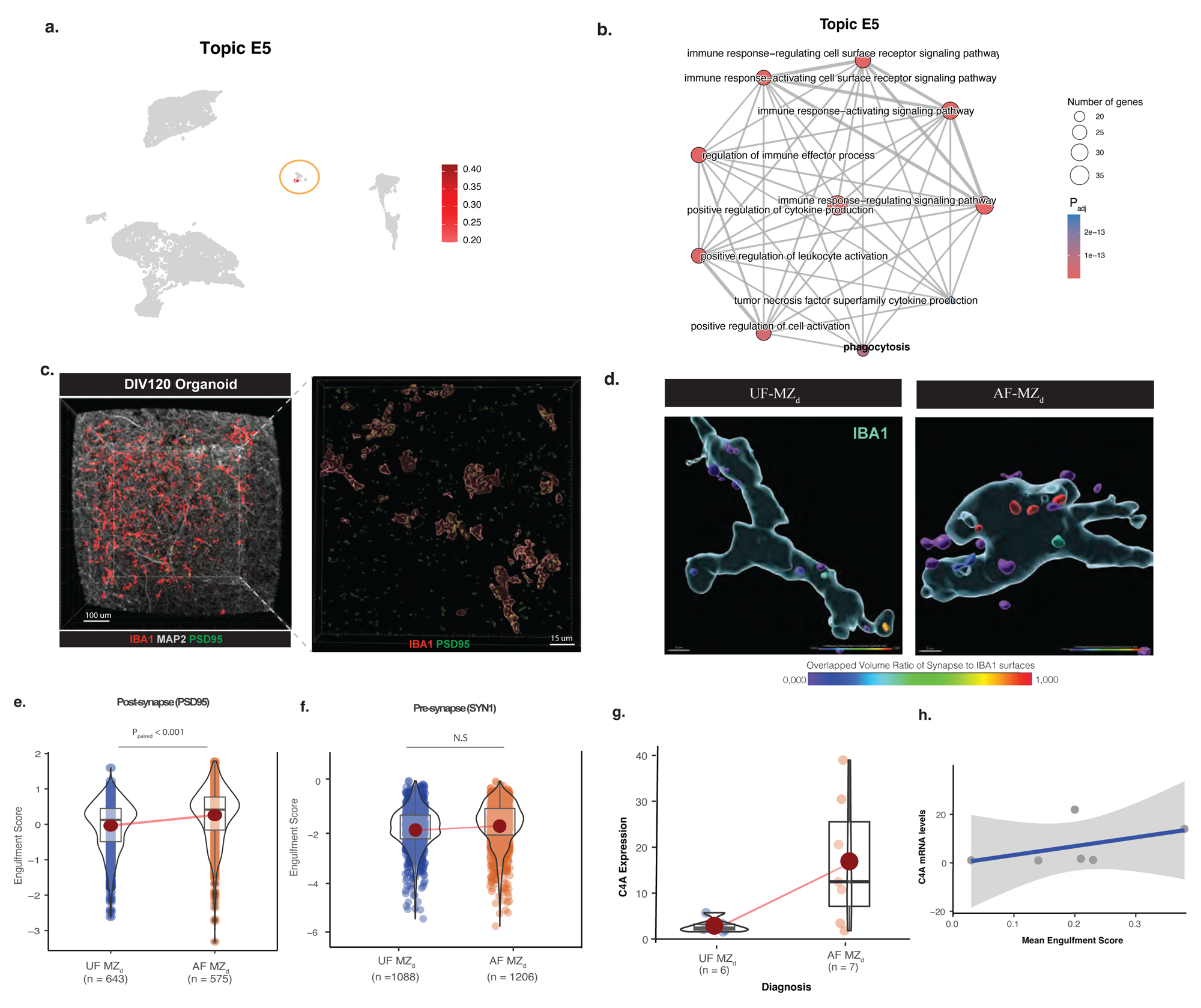
**(A)** UMAP of expression topic 5 (Topic e5) activation across microglia cells. Scale bar represents topic activation scores. **(B)** Enriched biological pathways (GO:BP Terms; P-adjusted<0.05) in Topic e5. Scale bar represents adjusted P values. Circle size represent number of genes mapped per term. **(C)** Confocal microscopy images of IBA1+ (red), MAP2+, and PSD95+ (green) stainings within DIV 120 organoids. A representative of the 3D reconstructions of the acquired images is shown on the right with IBA1+ cell bodies in red and PSD95+ synapse structures (overlapping: yellow and non-overlapping: green). **(D)** 3D reconstructions of confocal microscopy images of IBA1+ cell body (cyan) and PSD95+ structures (small dots colored by the ratio of overlapping volume with the cell body) from MZ_d_-derived organoids (DIV 120). Colour scale represents overlapping volume ratio of the synaptic structure and the microglial structure ensuring qualifying of synapses that are present only inside the cell body as engulfed. Quantification of engulfment of **(E)** post-synaptic structures (PSD95+), and **(F)** pre-synaptic structures (SYN1+) by microglia (IBA1+). Data points represent IBA1+ structures with corresponding engulfment scores ( overlapping volume ratio to synaptic structures; y-axis). Values are square-root transformed for visualisation. Significance is tested using linear regression models (Methods). **(G)** Violin plots of C4A mRNA expression levels via qPCR (estimate= 13.952, std.error= 5.540, t=2.518, *P*=0.02884) [C4B expression: estimate= 5.357, std.error= 4.199, t= 1.276, *P*= 0.2581]. Data points represent organoid replicates. Red dot indicates the median. Lower and upper hinges of the boxplot represent first and third quartiles. **(H)** Scatter plot with mean microglial engulfment of a subject (y-axis) and mean C4A mRNA expression of the subject (x-axis). Blue line represent linear fit with grey bars representing the 95% confidence interval.

Thus, we employed high-resolution microscopy and three-dimensional cellular reconstruction of microglia in DIV 120 organoids (Fig. 4c). As hypothesized, we observed a significant increase in microglial internalization of post-synaptic material (PSD-95) in organoids derived from SCZ-affected twins compared to their unaffected co-twins (mixed-effects model: estimate = 0.024, standard error = 0.007, t = 3.531; Methods; Fig. 4d–f).

Previous research has shown that microglial synaptic engulfment is mediated by classical complement signaling^40^, with structural variants in the *C4* gene—leading to increased expression of *C4A*, a known genetic risk factor for SCZ^41^—contributing to enhancing the microglial uptake of synaptic material^17,42^. To assess this, we measured *C4A* mRNA expression in the twin pairs using qPCR (n = 6 subjects). We observed elevated expression of *C4A* in SCZ-affected twins, while *C4B* expression—unrelated to SCZ risk—remained unchanged (Fig. 4g). Finally, we detected a positive, albeit statistically non-significant, correlation between microglial engulfment and *C4A* expression levels in DIV 120 organoids (Fig. 4h).

### Disease-associated astrocytic states

While there were no vast cell-type compositional differences, we observed subtle shifts in astroglia cell states. SCZ likelihood scores identified astrocytic subclusters (scz-AST1/2) enriched in SCZ, characterized by active TF regulomes and signatures targeting ciliary axonemes (*CFAP61, DNAH11*) components involved in cell cycle regulation during astrogenesis^43^, as well as genes linked to neuronal projection guidance (*SEMA5A, EPHA5, CNTN1, CNTNAP2*) within their inferred GRNs, albeit driven by one subject (Fig. 5a-e, Extended Data Fig. 6d-h).

**Figure 5.**
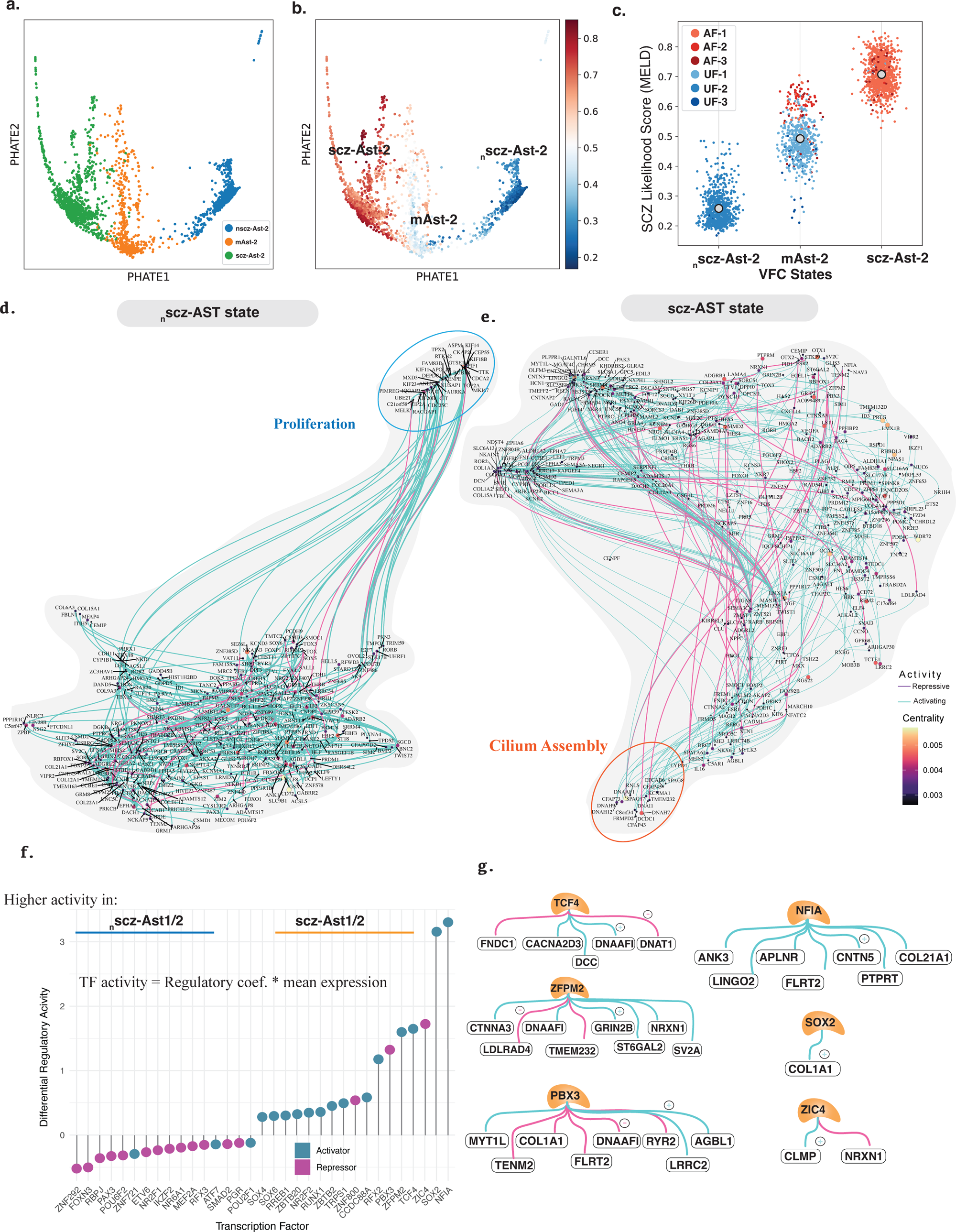
**(A)** PHATE embedding of Astrocyte-2 (AST-2) cluster with cells colored by the identified vertex frequency subclusters and **(B)** SCZ-likelihood scores. **(C)** Jitter plot (right) displays cells belonging to each MZ_d_ subject across both AST-2 cell states (x-axis) and their SCZ-likelihood scores (y-axis). AST-2 cell state enriched in SCZ-affected MZ_d_ is labelled as scz-AST-2 and unaffected MZ_d_ is labelled as _n_scz-AST-2. State that exhibits cells from both conditions is labelled as mAST-2. Legend indicates the MZ_d_ twin subject. **(D)** Gene regulatory networks (GRN; RNA+ATAC) of the SCZ-enriched AST-2 cell state (right), and unaffected MZ_d_ -enriched AST-2 cell state (left) computed by fitting generalized linear regression models for transcription factors (TFs), TF-binding motif accessibility, and expression of target genes, implemented in Pando. The TF networks are visualized on a UMAP embedding based on coexpression and inferred interaction strength between TFs. Highlighted parts TFs and target genes, involved in proliferation (blue) in unaffected-enriched astrocytic state, and cilium assembly (orange) in SCZ-enriched astrocytic state. Color and size of the dot represent PageRank centrality values of each significant TF (node; P-adjusted<0.05). Edges are colored by activity of the TF inferred by the sign of the regulatory coefficient (repressive-pink, activating-cyan). All significant TF-accessible peak-target gene trios of the GRNs are provided in Supplementary Table 10. **(F)** Top TFs in combined in astrocytes with differential activity between the SCZ-enriched cell states from both Astrocyte-1 and Astrocyte-2 clusters (scz-AST1/2). Activity for each TF is calculated as mean regulatory coefficient*mean expression of the TF. Differences in activities of each TF is shown on the y-axis with TFs on the x-axis. **(G)** Target genes of TFs with differential activity in astrocytes (scz-AST1/2 versus _n_scz-AST1/2), involved in cilium movement mechanisms. Cyan shows activation (+) and magenta shows repressive activity of TF on the corresponding target gene (−) as indicated by the regulatory coefficient.

In contrast, SCZ-depleted astrocyte states (_n_scz-AST) were defined by TF regulomes associated with mitotic processes, including *MKI67*, *TOP2A*, *NUSAP1*, and *KIF20B*—which were notably absent in the GRNs of scz-AST (Fig. 5d–e; Supplementary Table 10). Key regulatory modules distinguishing these networks included *NFIA*, *SOX2*, *ZFPM2*, *TCF4*, *ZIC4*, and *PBX3* (Fig. 5f), with target genes, including *NRXN1, GRIN2B, DCC* and *MYT1L*, thus enriching for known SCZ-risk variants and implicating trans-synaptic signalling and synapse assembly, similar to mechanisms indicated in the differential trajectory analysis (Fig. 5g; Supplementary table 11).

### Disruption in synaptic programs emerges early in neurodevelopment

Recent evidence suggests that rare and common risk variants for SCZ partly overlap^44^. Since our model was primarily designed to capture effects of rare SCZ *de novo* mutations^45,46^, we compared SCZ trajectory-associated genes with the latest set of common risk genes as identified by genome-wide association studies^8^. This revealed a significant overlap (*P* =1.1 × 10^−3^; Supplementary Table 12), suggesting that common risk variants may influence developmental trajectories in a similar manner.

We also compared the identified DEGs in our model to SCZ-signatures reported in postmortem SCZ brain studies^47–49^. Notably, DEGs from the developing neuronal clusters in our system did not significantly overlap with broadly defined neuronal DEGs from postmortem tissues (*P* = 0.084^47^ and *P* = 0.11^48^; Supplementary table 13), likely reflecting differences in developmental stage and disease progression. However, when focusing specifically on synaptic gene programs identified in postmortem studies^49^, we observed significant enrichment (*P* =2.8×10^−07^; Supplementary table 13). This suggests that disruptions in synaptic programs may emerge early during neurodevelopment and persist throughout the course of illness.

## DISCUSSION

In this study, we provide a unique (twin-based) single-cell multiome resource for understanding the intricate mechanisms underlying gene expression and chromatin accessibility of different cell types in the developing human brain in the context of SCZ. Using fate probability maps, we identify premature lineage commitment emerging from a distinct SCZ-associated radial glia cell state and reveal unique interactions among SCZ risk genes through unbiased multimodal analyses. The overall cell type composition for differentiated lineages remained largely unchanged but included SCZ-enriched subpopulations implicating perturbations in synaptic programs (neurons), trans-synaptic signalling (astrocytes), and phagocytosis (microglia). Consistent with these findings, and in line with the decreased synapse density in SCZ patients, we also observed an increased uptake of synaptic material in microglia.

Fine mapping studies of SCZ high-risk variants, combined with temporal gene expression patterns of implicated risk genes, strongly suggest that genetics-driven disease liability is likely to manifest already during the second trimester, i.e., the developmental time-window modeled in this study. Notably, SCZ trajectory-associated genes were enriched for both common and rare genetic risk variants for SCZ. However, these genes did not significantly overlap with DEGs from SCZ postmortem brain tissue. For instance, the observed interaction between *ZEB2* and *SORCS3* within the SCZ-enriched radial glia subpopulation underscores the value of modeling early brain development, and reinforces the idea that brain organoids can offer additional and important insights into the functional impact of SCZ risk factors.

A shift in the developmental timing of neuronal birth in the telencephalon is critical to subtype-and cortical layer-specification. Thus neurons born prematurely are likely to exhibit dysregulated migratory cues, facilitating abnormal connectivity, circuit formation and plasticity^50–52^. Upon leveraging the two modalities, we were able to map biological networks and infer key regulators that drive progenitors towards an abberant state, which have been shown to influence neuronal subtypes, e.g., specification of upper-layer neurons implicated in SCZ^47^. SCZ-enriched neuronal subpopulations showed disrupted synaptic signaling, with enrichment of synaptic programs linked to genetic risk variants. Similarly, SCZ-enriched astrocytic subpopulations exhibited perturbations in synapse assembly and trans-synaptic signalling. Together, these disruptions likely contribute to the increased uptake of synaptic material in microglia as shown in this study, which is likely mediated by upregulation of microglial phagocytic genes due to changes in the neural environment rather than primarily being an effect of intrinsic microglial factors.

Two key technical aspects of this study are the use of the rare monozygotic twins discordant for SCZ and the application of a single-cell multiome approach. Although the twin design comes at the expense of a smaller sample size, the proteomic data highlighted similarities according to diagnosis rather than twin pair, and the genetic enrichment analysis based on multiomic data captured established SCZ risk genes, thereby confirming that our model captures risk factors sufficient to drive phenotypic differences despite the twins’ nearly identical genetic background. The strength of the paired multiome approach also includes the ability to cross-validate gene expression changes to the accessibility of TFs and the outcome of their binding activity on target genes at a single cell resolution. This consistency across modalities thus enhances statistical power and the likelihood of deciphering true biological associations by reducing additional noise commonly present in next generation sequencing data.

Finally, our findings are restricted to the fetal phase of brain development as brain organoid models recapitulate early developmental phenotypes. Future studies aimed at investigating downstream effects of these early alterations at later developmental points may provide better insights into mechanism underlying the emergence of the symptomatic spectra in SCZ.

## ONLINE METHODS

### Ethics and iPSC reprogramming

Three pairs of monozygotic twins discordant for schizophrenia were recruited in Finland^53^. Subject-specific demographics are provided in Supplementary Table 1. Each subject was diagnosed by a trained psychiatrist according to DSM-IV criteria and assessed using the Positive and Negative Syndrome Scale (PANSS) at the time of sampling. All study subjects provided an informed consent. Approval of this study has been granted by the Ethics Committee of Helsinki University Hospital District (HUS/1391/2016), and Swedish Ethical Review Authority (#2023-05308-01).

Skin biopsies were obtained to derive fibroblasts which were expanded in a fibroblast culture media containing DMEM, 20% fetal bovine serum, 1% Penicillin-Streptomycin, and 1% non-essential amino acids. Fibroblast cultures of all subjects tested negative for mycoplasma contamination using the LookOut Mycoplasma PCR detection kit (Sigma Aldrich, MP0035). Fibroblasts were then reprogrammed into iPSCs using the CytoTune-iPS 2.0 Sendai reprogramming kit (ThermoFisher Scientific) according to manufacturer’s instructions, involving transduction with three vectors carrying *hOCT3/4*, *hSOX2*, *hKLF4* and *hc-MYC* to induce pluripotency. All iPSCs were expanded in mTeSR Plus Medium (STEMCELL Technologies, # 100-0276) on GelTrex-coated plates and purified using magnetic cell sorting (MACS) with Anti-TRA 1-60 beads (Miltenyi Biotech, 130-100-082) according to manufacturer’s instructions. All iPSC cultures were negative for mycoplasma, and stained positive for OCT4 via immunocytochemistry, confirming pluripotency.

### STAR Cohort

The schizophrenia and bipolar twin study in Sweden (STAR) includes MZ and DZ twin pairs with schizophrenia or bipolar disorder, and a total of 462 twins have participated, see Johansson et al. for further cohort description^54^. Study participants were identified through the Swedish Twin register and the National Patient register, and they were invited to join the STAR study if only one twin had a registered episode of schizophrenia or bipolar disorder. Written informed consent was obtained from all participants in this study. Ethical permissions were obtained from Stockholm County in Sweden (Dnr: 2004/448/4, 2007/779-31/3 and 2008/292-32.

Participating twins received extensive assessment procedures. Diagnostic status was confirmed by a clinical psychiatrist through the Structured Clinical Interview for DSM-IV (SCID-I), and the final diagnosis was determined by an evaluation team, incorporating data from previous hospitalizations and hospital records. The diagnoses were categorized as schizophrenia (ICD-10: F20), schizoaffective disorder (ICD-10: F25), bipolar disorder (ICD-10: F31), major depressive disorder (ICD-10 F32-F33) or not affected by any of those diagnoses. From the STAR cohort, 19 disease discordant MZ twin pairs were included for whole genome sequencing in which one twin was diagnosed with schizophrenia, schizoaffective disorder, or bipolar disorder, and the co-twin was not diagnosed with any of these disorders^55^. While major depressive disorder (MDD) can involve psychosis, we included pairs where the co-twin had a diagnosis of MDD without occurrence of psychotic symptoms and considered that subject as unaffected. For the final analyses shown in this study, 9 MZ twin pairs, diagnosed with schizophrenia and schizoaffective disorder only, were included.

### WGS and CNV calling

The procedures followed for WGS and subsequent CNV calling have been previously documented^55^. Briefly, blood samples were collected during clinical ascertainments and DNA extraction was performed. One twin pair was excluded from WGS after failing DNA quality assessment by agarose gel electrophoresis. WGS was performed by Edinburgh Genomics (Clinical Genomics) on a HiSeqX to an average depth of coverage of 30x per sample. Reads were aligned to the GRCh38 reference genome using BWA-MEM^56^, following the GATK Best Practices^57^.

Four separate calling tools were used to define the germline CNVs. All the calls within a twin pair were combined to create Regions of Interest (ROIs). Any ROI found in only one twin and not their co-twin, and identified by only one calling algorithm, was removed. Variants with at least 50% reciprocal overlap with common variants (>1% frequency) in the appropriate reference population were removed unless it was labelled “Pathogenic”in the NIH Clinical Genomics (ClinGen) CNV database (UCSC “iscaPathogenic” table)^58^.

After quality control, 17 twin pairs remained for analysis, and principal component analysis determined that all were of European ancestry except one twin pair from East Asia. Comparison with existing genotype array data confirmed expected sample identities and zygosities.

### Generation of microglia-containing brain organoid cultures

Microglia-containing brain organoids were generated according to a previously published protocol with a few modifications^18^. Briefly, human primitive neural precursor cells (pNPCs) and human primitive macrophage progenitors (PMPs) were both generated from respective patient-derived iPSC lines. Human PMPs were derived in a Myb-independent manner by seeding 10,000 iPSCs per well on ultra-low attachment V bottom 96 well plates, cultured in mTeSR plus medium supplemented with rock inhibitor (10uM; Sigma Y0503) for 24 hours. The media was carefully changed to pre-PMP embryoid body (EB) medium containing mTeSR plus media supplemented with BMP-4 (50 ng/ml), SCF (20 ng/ml), VEGF (50 ng/ml) and Y-27632 rock inhibitor (10uM). The emerging EBs were observed for 4 days and transferred to a tissue culture treated 6-well plates in PMP media containing X-VIVO15 supplemented with M-CSF (100 ng/ml), IL-3 (25 ng/ml), GlutaMAX (1X), penicillin-streptomycin (P/S; 1X), and β-mercaptoethanol (100 uM). The media was changed once every week for approximately three to four weeks, when free-floating PMP cells started to appear in suspension which were harvested and co-cultured with pNPCs.

For generation of pNPCs, we seeded 4000 iPSCs on V-bottom ultra-low-attachment 96-well plate and cultured for 1 week in EB medium containing Advanced DMEM/F12 supplemented with dorsopmorphin (1 uM), N-2 (1X), B27 -RA (1X), and SB431542 (5uM). EBs were then transferred to growth factor reduced Matrigel-coated 6-well plates and cultured for 1 week in media containing DMEM/F12 supplemented with N-2 (1X) and Laminin 521 (1 ug/ml) to induce neural rosettes. After 7 days, the rosettes were manually isolated and expanded for 1 week on growth factor reduced Matrigel-coated 6-well plates in NPC media composed of Neurobasal and Advanced DMEM/F12 (1:1), supplemented with N-2 (1X), B27 -RA (1X), FGF2 (20 ng/ml), hLIF (10 ng/ml), CHIR99021 (3 uM), SB431541 (2 uM) and Y-27632 rock inhibitor (10 uM). The NPCs were purified using MACs sorting for CD27-negative fraction and CD133-positive fraction according to manufacturer’s instructions.

3000 PMP cells and 7000 NPC cells derived from each iPSC line were then combined per organoid in a 7:3 ratio, plated per well in an ultra-low attachment 96 well plate and centrifuged at 150xg for 5 minutes at RT. The resulting EBs were transferred to an ultra-low attachment 24 well plate and incubated at 37°C in proliferation medium, containing 1:1 mixture of NPC and PMP media for DIV 4, with fresh media changes every day. At DIV 27, the plates were moved onto an orbital shaker set at a speed of 80-90 rpm within the incubator. At DIV 35, the organoids were transferred to ultra-low attachment 6-well plates and the media was changed to a maturation media containing DMEM/F12, Neurobasal and BrainPhys neuronal (2:1:1) media, supplemented with N-2 (1X), BDNF (20 ng/ml), GDNF (20 ng/ml), dibutyryl cyclic AMP (1 uM), IL-34 (100 ng/ml), GM-CSF (10 ng/ml), ascorbic acid (200 nM), and P/S (0.5X). The media was changed every other day until DIV 120 and harvested for further experiments.

### Organoid harvesting and Immunohistochemistry

Organoids were washed in PBS (1X) and fixed in 4% PFA for 45 minutes at RT followed by two PBS washes and incubated in 30% sucrose solution (w/v) overnight at 4°C. The organoids were then embedded in OCT (HistoLab, #45830) and flash frozen using dry ice and isopropanol and stored at −80°C. For immunohistochemistry, frozen organoid blocks were cryosectioned using a NX70 cryostat equilibrated at −18°C with a blade temperature of −16°C to obtain 18-um thick organoid sections at 20 um intervals. The cryosections were washed with PBS (1X) and exposed to permeabilization and blocking solution containing 0.3% Triton X-100 in PBS (1X) with 3% BSA, for 60 minutes at RT. The sections were washed with PBS (1X) and incubated overnight at 4°C in primary antibodies diluted in the blocking solution. The sections were washed thrice in PBS (1X) for 5 minutes each followed by incubation with secondary antibodies diluted in the blocking solution for 2 hours at RT in the dark. The sections were then washed thrice in PBS (1X) and incubated with DAPI (1:500) for 5 minutes at RT before mounting on glass slides in a fluorescence mounting media (DAKO, #S302380-2).

For whole-mount organoid immunostainings, organoids were harvested and fixed in 4% PFA for 1-3 hours depending on their size. The organoids were washed twice in PTx.2 mixture (2% Triton X-100 in PBS) for one hour each at 37°C on an orbital shaker. The organoids were then incubated overnight on a shaker at 37°C in permeabilization solution (PTx.2 supplemented with 0.02% glycine (w/v) and 20% DMSO). The next day the organoids were incubated in blocking solution (PTx.2 supplemented with 6% BSA and 10% DMSO) for 6 hours followed by overnight incubation with primary antibody solution (2X) diluted in PTwH mixture (2% tween-20, 10ug/ml Heparin in PBS) at 37°C on a shaker. The organoids were washed 4 times in PTwH mixture for 2 hours each and incubated overnight with secondary antibody solution (2X) diluted in PTwH mixture at 37°C on a shaker in the dark. The organoids were washed 4 times in PTwH mixture for 2 hours each and incubated with DAPI (1:500) for 2 hours at RT in the dark and washed with PTwH for 1 hour at RT. Optical clearing was performed on the organoids using the iDISCO+ method^59^. The immunostained whole organoids were embedded in 1.5% agarose and dehydrated in a 4-step methanol series (20%, 60%, 80% and 100%) for 1 hour each. The organoid blocks were then incubated in 1 volume of Methanol/2 Volumes of Dichloromethane (DCM) solution for 30 minutes, followed by two washes in 100% DCM, 20 minutes each, and transferred to DiBenzyl ether (DBE, Sigma 1080141KG) until clear and stored in DBE at RT in the dark.

### Image acquisition and quantification

Immuno-stained organoid sections were imaged on ZEISS LSM900, an inverted laser scanning confocal microscope with an Airy-scan 2 detector for super-resolution imaging. Images were acquired with 2 objectives, 40X water (NA=1.2, WD=0.29mm, C-Apochromat), and 20X air (NA=0.8, WD=0.55mm, Plan-Apochromat). 405nm, 488nm, 561nm and 640nm lasers were used for acquisition. Whole-mount images of cleared organoids were acquired on ZEISS LSM800 confocal microscope using custom 3D printed chambers to hold the organoid blocks in DBE and imaged using a 10X air objective (NA=0.3, WD=5.2mm, EC Plan-Neofluar). Entire organoid images were obtained via tile acquisition on the ZEN2009 software.

The raw images (czi format) with Z-stacks (18um range; 1um steps) were processed using the Imaris software (v0.1; https://imaris.oxinst.com/). Three dimensional volumetric reconstructions of channels capturing IBA1 and PSD95 markers were performed to obtain microglial cell surfaces and synaptic structure surfaces respectively. Background subtraction was performed followed by smooth object reconstruction. Each object in the IBA1 channel was defined as a microglial cell and each object in the PSD95 channel was defined as a synaptic structure. Synaptic structure surfaces with the overlapping volume ratio with microglia > 0.9 were defined as internalized synaptic surfaces. To quantify microglial engulfment of synaptic structures, colocalization of IBA1 and internalized PSD95 surfaces was quantified by calculating the ‘Overlapping volume ratio between the IBA1 and PSD95 surfaces’ (engulfment index), and the ‘Shortest distance to PSD95 surfaces’ per microglia cell, normalized by the volume of cellular surfaces. The representative images depict the synaptic structures statistically color-coded, ranging from 0 to 1, according to the overlapped volume ratio with the microglial cell surface, indicating engulfment.

### Electrophysiology

Coverslips were placed in a submerged chamber perfused with aerated artificial cerebrospinal fluid (ACSF; in mM: 124 NaCl, 30 NaHCO3, 10 glucose, 1.25 NaH2PO4, 3.5 KCl, 1.5 MgCl2, 1.5 CaCl2), at 36°C with a perfusion rate of 1-2 ml per minute. Coverslips were left undisturbed for 5 minutes before any recording. Patch-clamp (whole-cell) recordings were performed with borosilicate glass microelectrodes (4–6 MΩ) from the soma of visually identified cells using IR-DIC microscopy (Zeiss Axioskop, Germany) using a Multiclamp 700B (Molecular Devices, CA, USA). For neuronal electrophysiological properties (Vh = −70 mV) a potassium-based intracellular solution was used (in mM): 122.5 K-gluconate, 8 KCl, 4 Na2-ATP, 0.3 Na2-GTP, 10 HEPES, 0.2 EGTA, 2 MgCl2, 10 Na2-Phosphocreatine, set to pH 7.2-7.3 with KOH, osmolarity 270-280 mOsm. The signals were sampled at 10 kHz, software low-pass filtered at 2 kHz, digitized and stored using a Digidata 1440A and pCLAMP 10.4 software (Molecular Devices, CA, USA).

### Quantitative PCR (qPCR)

Expression levels of C4, C4A and C4B mRNA was measured using qPCR. 3 whole organoid lysates per line were harvested in Tri reagent (Sigma, #T9424) and processed for RNA extraction using the DirectZol RNA-Miniprep Kit (Zymo Research Inc., #R2050), according to the manufacturers protocol. DNAse I treatment was performed for on bound RNA in Zymo-spin columns. The RNA was then washed and eluted in 30μl nuclease-free water. Quality of the eluted RNA was determined using NanoDrop (Thermofisher Scientific) and stored at −80°C.

The total RNA was thawed at 4°C and reverse transcribed to cDNA using the High-Capacity RNA-to-cDNA Synthesis Kit (Thermofisher Scientific) according to the manufacturer’s instructions, in 20μl reaction volumes containing 1μg of input RNA, on a thermal cycler. The cDNA was further diluted to 1:2 and used as the input for the qPCR reactions. qPCR was performed using the TaqMan Gene Expression Assay (Applied Biosystems) on StepOnePlus RealTime PCR system (Applied Biosystems). Reactions were carried out in 20ul volumes containing 5μl cDNA, 300 nM forward and reverse primer, 250 nM probe concentration, and 10μl TaqMan PCR master mix (Applied Biosystems, #4324018). The primer and probe sequences used were as follows:

Forward Primer: GCAGGAGACATCTAACTGGCTTCT
Reverse Primer: CCGCACCTGCATGCTCCT
*C4A*-probe: FAM-ACC CCT GTC CAG TGT TAG-MGB
*C4B*-probe: FAM-ACC TCT CTC CAG TGA TAC-MGB

Relative expression levels of *C4A* and *C4B* gene were normalized against the housekeeping genes, *GAPDH* (Assay ID. Hs99999905_m1, Thermofisher Scientific), and quantified according to the ΔΔC_t_ method. Fold changes (2^−ΔΔC^ ) were calculated by normalizing ΔC_t_-values of each sample to the average ΔCt-value of each subject. Differences in expression levels across condition were tested using a linear model, accounting for twin pairs (Fold change ∼ condition + pair).

### Single nucleus Multiomic Analyses

#### Nuclei Isolation

For each subject from the three twin pairs, six fresh organoids were pooled at DIV 120, mechanically dissociated on ice into a single cell suspension in NbActiv1 media (Neurobasal medium, B27, and GlutaMax supplement) and triturated with glass capillary pipettes of decreasing inner diameters to obtain a homogenous suspension. The suspension was filtered through a 70 micron filter and centrifuged at 500 RCF for 5 minutes at 4°C. Nuclei isolation was performed according to the Chromium Next GEM Single Cell Multiome ATAC + Gene Expression (GEX) protocol (CGOOO338). The supernatant was discarded, and the cell pellet was incubated in 100μl of chilled 0.1X Lysis buffer for 8 minutes at 4°C (*the lysis time was optimized for maximal cell lysis and minimal nuclei overlysis*). The lysed cellular mixture was diluted with 900μl of wash buffer and filtered through a 37micron filter, followed by centrifugation at 500 RCF for 5 minutes at 4°C. The obtained pellet was washed and subjected to iodoxinol gradient (OptiPrep™ Sigma-aldrich; 50% and 29% Iodixanol in Iodixanol medium containing 2M sucrose, 2M KCl, 1M MgCl2 and 1M Tris buffer) to get rid of cellular debris and centrifuged at 13500 RCF for 20 minutes at 4°C. Pure nuclei pellet was carefully isolated and centrifuged at 500 RCF for 5min at 4°C in wash buffer before resuspending the final nuclei pellet in Diluted Nuclei buffer (containing Sigma-Aldrich Protector RNAse inhibitor) to obtain a desired nuclei concentration of 1700 nuclei/μl. The quality of nuclei was assessed under a microscope at 40X and counted using a *Berker Chamber* prior to proceeding with Chromium Next GEM Single Cell Multiome ATAC + Gene Expression (CG000338) Rev E protocol. The nuclei suspensions were incubated in a transposition mix followed by GEM generation and barcoding of full-length cDNA from polyA mRNA for gene expression libraries and transposed DNA fragments for ATAC libraries. One library per subject per modality was generated. The libraries were quantified using the High sensitivity DNA Assay (Qiagen) and fragments were assessed on a Bioanalyzer (Agilent technologies) prior to pooling for sequencing on Illumina NovaSeq S6000 (S2-100 v1.5 flowcell).

#### snMultiome Data Analysis

Raw single nuclei multiome GEX and ATAC sequencing data (fastq) were processed using CellRanger ARC (v1.0,10X Genomics). For both modalities individually, alignment (human; GRCh38), filtering, peak calling and counting was performed and aggregated to obtain filtered UMI count and binary peak barcode matrices. Gene expression counts were filtered for ambient RNA contamination using CellBender^60^ and ATAC fragments file were used to call peaks using MACS2^61^. The resulting data was pre-processed using Seurat (v4)^62^ and Signac (v1.1)^63^ with paired gene expression and chromatin accessibility profiles per cell from all samples. Quality control was performed on each modality per-cell with high quality cells containing 1000 < UMI count < 50000, 1000 < RNA features < 12000, mitochondrial percentage < 5%, 1000 < ATAC count < 100000, 1000 < ATAC features < 30000, nucleosome signal < 2, TSS enrichment score > 1.

#### Topic Modeling

To integrate gene expression and chromatin accessibility, we leveraged MIRA^64^, a recently developed tool that employs joint probabilistic modelling of multimodal data using variational inference and Bayesian inference to capture dependencies between both datasets. We employed the CODAL algorithm implemented in MIRA for batch correction to remove unwanted batch effects across 6 batches. An anndata object for each modality was generated and gene expression was log-normalized in Scanpy. To increase the training speed, 2500 highly variable genes (dispersion threshold= −0.1) were used as encoder features to initialize the expression topic model while all accessibility peaks were used as encoder features to initialize the accessibility topic model. We used the learning rate range test to determine learning rate boundaries (RNA: 10^−3^ – 10^−1^; ATAC: 10^−5^ −10^−1^) to ensure the best fit and proceeded to determine the appropriate number of topics by Bayesian hyperparameter optimization per modality. Over 50 iterations (best score = 2.28×10^3^), we obtained 19 and 24 topics, trained the expression or accessibility topic models with the tuned hyperparameters. The trained expression and accesibility topic models were used to predict cell-topic compositions and integrated the transformed topic spaces with a box-cox transformation parameter of 0.25, to construct a joint KNN graph visualized using UMAP. To evaluate how expression and accessibility topics correspond to each other, we calculated pointwise mutual information between RNA and ATAC topic compositions per cell. For expression topics, functional enrichments of all genes within topics were performed whereas for accessiblity topics, peaks within each topic were scanned for transcription factor binding sites and annotated as TF motifs using the hg38 genome and JASPAR 2022 database. Top 10% peak quantile representing each topic were used to retrieve motif enrichments for each topic. Motif scores were calculated using get_motif_scores function in MIRA.

To assess association of each topic to diagnosis status (SCZ-affected MZ or unaffected MZ), we compared topic activation scores per cell across groups by fitting a logistic regression model of individual topic values to diagnosis, adjusted for twin pair. The condition associated with higher cell-topic scores were based on the sign of the estimated coefficient. Topics with positive coefficients with *P <* 0.05 were determined as SCZ-affected topics whereas topics with negative coefficients and p<0.05 were associated to unaffected twins. [glm(Condition ∼ Topic + TwinPair, family = binomial)]. In case of continuous variables, such as estimating association of topics to computed pseudotime values, a linear regression was performed for each topic with an interaction term between pseudotime and group, adjusted for twin pair. [glm(Topic ∼ Pseudotime * Group + TwinPair)]. Condition associated with higher cell-topic scores were based on the sign of the estimated coefficient for the interaction term.

#### Reference mapping and Cell type annotation

Cluster-specific gene expression signatures were computed by performing differential expression analysis in Scanpy (rank_gene_groups) using wilcoxon rank-sum tests on the identified clusters. Fasta sequences from the human hg38 were downloaded from the UCSC repository and scanned for motifs to find TF binding sites in peaks using the get_motif_hits_in_peaks function implemented in Mira. TFs with motifs without expression in the RNA modality were filtered out. Cell type-specific motif activity was calculated similar to gene expression. Multimodal reference mapping for the entire dataset was performed using label transfer techniques implemented in Seurat with a multiome dataset (gene expression and chromatin accessibility) from primary fetal cortex (Zhu et al., 2023). The dataset was normalized using SCTransform and anchors (PCA) between the query and reference dataset were computed using FindTransferAnchors. Cells from the reference dataset were projected onto UMAP model used earlier on the joint KNN graph using MapQuery. After mapping, cell type labels from the fetal cortical dataset were transferred onto the organoid dataset. The projection scores of individual labels were visualized per cell across the clusters in the UMAP embedding. Cell type annotations of the organoid dataset were confirmed after comparing transcriptomic correlations (pearson’s correlation) to scRNAseq datasets from the developing fetal brain across first, second, and third trimester, followed by postnatal ages, as well as the adult human brain, using clustifyR package (2000 variable features). Bulk RNA-seq data from BrainSpan atlas of the human developing brain was also compared to aggregated transcriptomes per cluster and the entire organoid to match an overall developmental period to the model.

#### Differential composition testing

Shifts in the composition of cell types within the dataset was performed for all identified clusters using compositional data (CoDa) framework of cacoa R package^65^ designed for case-control comparisons of scRNAseq data. The RNA modality was pre-processed using Seurat and a cacoa object was initiated with twin subjects as the sample groups. CoDa loadings were visualized and statistical significance was estimated at a Bonferroni threshold of 0.01.

#### Pseudo bulk differential analyses and pathway enrichment

To identify modality specific dysregulated elements, pseudo-bulk differential analyses were performed by selecting 100 cells per subject per group at random in every cell type. Raw expression or peak counts were aggregated into ‘bulk’ samples and processed using DESeq2 method (likelihood ratio test) and fitting gamma-poisson generalized linear models^66^ on raw counts with design = ∼ twin pair + condition to obtain differentially expressed genes (DEGs) or differentially accessible region (DARs). Genes linked to DARs were inferred by association using GREAT functional enrichment analysis^67^. Pathway analysis was performed downstream of pseudo bulk differential analysis by ranking genes according to the statistic and performing fast gene set enrichment analysis using fgsea R package^68^ to obtain normalized enrichment scores for pathways in the Gene Ontology: Biological pathways database. To obtain functional enrichment in shortlisted set of genes or regions, pathways were computed using overrepresentation analysis implemented in gprofiler R package. Cellular crosstalk via ligand-receptor inference between cell types was performed using CellChat tool^69^. Ligand-receptor (LR) signalling was determined by computing communication probabilities mediated by expression of ligand and receptors from precursors and glia to neurons. Differential expression analysis of signalling was performed by comparing fold change of ligands in secreting cell type and receptors in the receiving cell type between the two conditions per sender-receiver cell types. Significance is tested using paired Wilcoxon tests at p-value < 0.05.

#### Trajectory inference and fate probability

The undirected joint KNN-graph was used to compute and assign pseudotime values to each cell implemented in Mira. A principal tree was fit on the pseudotime-arranged cells using scFates package^70^ to identify bifurcations and uncover existing trajectories and a dendrogram of the tree were estimated for each condition separately. CellRank python package^71^ was used to compute a cell-cell transition matrix using the PseudotimeKernel and passed to a GPCCA estimator to identify terminal macrostates with radial glia as the initial state. Fate probabilities for each terminal state were calculated by aggregating all random walks towards a terminal state and compared per cell across subjects using Mann-Whitney U test (*P*<0.05). We analysed expression dynamics along trajectories in the joint KNN graph using tradeSeq R package. Generalized additive models (GAM) models were fitted for each gene as a smooth function of pseudotime using the fitGAM function separately on neuronal lineage cells (Radial glia, IPC, neuronal clusters) and the glial lineage (radial glia and astrocytic clusters) accounting for twin pairs as a covariate. The association to pseudotime and diagnosis was statistically tested using conditionTest in tradeSeq. Differential progression and differentiation of lineages were statistically tested using progressionTest (custom KS test, p<0.01) and DifferentiationTest (random forest classifier, p<0.05) functions implemented in condiments workflow. For visualizing, gaussian smoothing (sigma=1) was applied to the expression data per gene.

#### Disorder-associated cell states (MELD)

Cellular states associated to SCZ were determined by quantifying the likelihood of observing a particular cell state in either SCZ-affected twins or their unaffected co-twins, using sample-associated density estimates and relative disorder likelihoods per cell over a manifold of cell states with functions implemented in the MELD python package ^72^. Data was embedded using PHATE and a KNN graph was built (knn=10, decay=10) on normalized data. Optimal filter and graph parameter tuning were done using meld.Benchmarker to get knn=11, beta =54 and mean square error = 8.24×10^−4^. The output from the MELD operator assigned each cell to a SCZ likelihood score and heterogeneity in scores within clusters were examined using PHATE embedding of individual cell types. Cells from radial glia, astrocyte-1 and astrocyte-2 clusters displayed varying SCZ likelihood scores which were further subjected to vertex frequency clustering (VFC) implemented in meld.VertexFrequencyCluster to discover cellular states that have similar transcriptional responses in a particular condition.

#### Gene regulatory network inference and TF activity

To construct gene regulatory networks of disorder-associated cell states, we isolated the corresponding cells and applied the Pando framework individually^24^. A GRN object was initiated with all called peaks and SCT-normalized counts with computed variable features. TF-binding motifs were scanned across regions using curated motifs from JASPAR collection within Pando, to predict binding sites. 40,000 random peaks were chosen as a background peak list. The network is constructed using infer_grn, retaining gene-correlated peaks (0.05) estimated by Signac, and fitting generalized linear model per gene as a function of TF expression and its binding site accessibility. Model coefficients for each TF-peak-target trios and adjusted p values were extracted to keep significant interactions (*P_adj_*<0.01). Regulatory modules were identified using find_modules with an R^2^ threshold of 0.1. The network was visualized using the igraph package in a UMAP embedding. To compare drivers of differences between two cell states, TF activity was defined by multiplying the mean regulatory coefficient with the mean expression of a TF in each state and differences in mean activity of each TF between two states were computed. The type of activity (repressive or activating) was determined by the sign of the mean regulatory coefficient (positive-activating; negative-repressive).

#### Enrichment of SNP-based risk variants

SCZ summary statistics were downloaded from the latest SCZ GWAS publication and processed for gene associations using MAGMA software^73^. Results were compared to the final gene set provided by the authors for a consensus risk gene set for schizophrenia. Enrichment of schizophrenia risk genes in the identified trajectory-associated dysregulated genes from the neuronal and glial lineages was tested using one-sided Fishers exact test (P<0.05).

#### Post-mortem and SCZ-associated signature comparisons

Lists of disorder-specific gene signatures were downloaded from two SCZ postmortem studies with snRNAseq data^47,49^. For Batiuk et al., significant differentially expressed genes (P-adjusted < 0.05) for all neuronal clusters were combined and tested for overlap with differentially expressed genes of neuronal clusters of this study (CPN-1, CPN-2, CFuPN-1, CFuPN-2, Interneuron, uPN, nCPN-1, nCPN-2, nPN). For Ling et al., the pre-defined SNAP-a and SNAP-n program genes were obtained. Cells from astrocyte clusters of this study (Astrocyte-1, Astrocyte-2, Astrocyte-3) were scored for the SNAP-a program, and cells from neuronal clusters of this study were scored for the SNAP-n program using AddModuleScore function in Seurat. Both SNAP programs were also tested for overlaps in differentially expressed genes from cell type clusters of this study. All overlap comparisons were tested for significance using one-sided Fishers exact test (alternative=greater; *P*<0.05).

#### Mass spectrometry-based proteomics

Bulk proteomics on three organoids pooled per subject was performed using TMT-labelled mass spectrometry. The resulting data from TMT-based quantification in Proteome Discoverer using raw abundances contained 6 (samples) by 4999 (proteins) data matrix. Standard processing including log transformation, filtering missing peptide intensities was conducted using msqrob2 R package^74^. Peptides were aggregated and summarized into proteins and normalized data was visualized using principal component analysis (PCA) followed by hierarchical clustering with functions from factomineR package.

#### Genomic regions STAR cohort

Genomic coordinates of regions that were discordant in at least 4 MZ twin pairs were extracted and annotated using CHIPseeker R package. Genes mapped to these regions were tested for brain-specific expression during prenatal and postnatal brain development using the BrainSpan transcriptome dataset (www.brainspan.org; bulk RNA-seq). Co-expression networks were determined by calculating the pairwise Pearson’s correlation coefficient (R) between the seed gene (e.g. *SORCS3*) and 44319 features available in the BrainSpan dataset. *P*-values were adjusted for multiple testing using FDR. Positive networks included genes with R > 0 and FDR < 0.05 whereas negative networks included genes with R < 0 and FDR < 0.05. The co-expression networks were tested for biological pathway enrichment (Gene Ontology: BP) using clusterProfiler (*P_adj_* < 0.05).

#### Statistical Analysis

Three monozygotic twin pairs discordant for SCZ were investigated with three organoid replicates and two technical replicates per subject line were used for analyses involving imaging and qPCR quantification. Subjects within each group were paired according to the twin pair in all adjusted analyses. Comparisons across conditions were tested using a linear mixed effects regression model (grouping variable = twin pair with a random intercept) using the lme4 R package^75^. Covariates such as age, sex, replicates, and organoid sections (independent variables) were tested for confounding effects within the model (added as fixed effects). P-values for the linear models were generated using ANOVA and reported along with beta estimates, standard errors, and t-values. Within-twin pair comparisons were tested using the Wilcoxon signed-rank test (non-parametric). Normality assumptions were assessed where necessary. Correlations were tested using Pearson’s correlation. Statistical analyses for single nuclei multiomic and proteomic data are described in their respective methods sections. All statistical analyses were conducted using R (version 4.2.2) running under macOS (Ventura 13.6.1).

## DATA AVAILABILITY

Processed RNA and ATAC count matrices, along with cell metadata files required to replicate findings of this study, will be made available upon publication. Raw sequencing data is under controlled access due to applicable ethical approvals. Specifically, data could be available but for academic non-commercial research purposes only, and is subject to ethical review. Other source data can be provided upon request. Code used to process the single-nuclei multiomic and proteomic data presented in this study will be made available at https://github.com/SellgrenLab/SCZ-twins-multiome.

## ACKNOWLEDGEMENTS

We are grateful to to all the patients who participated in this study by donating their samples. We thank Åsa Björklund and Konstantin Khodosevich for their valuable suggestions and discussions contributing to this study. We also thank Peri Noori, Renuka Kudva, and Ákos Végvári for their help with library preparations, sequencing, and proteomic sample processing, respectively. We acknowledge Mukund Kabbe for his help with nuclei isolation. The computations/data handling was enabled by resources provided by the National Academic Infrastructure for Supercomputing in Sweden (NAISS), partially funded by the Swedish Research Council through grant agreement no. 2022-06725.

## CONTRIBUTIONS

C.M.S., S.M., J.G.L., and S. conceived the study. I.O. and O.V. diagnosed, assessed, and sampled the subjects. J.K., J.T., Š.L., and M.K. provided the iPSCs. J.G.L. and A.O.O. expanded the iPSCs and NPCs. J.G.L. and S.M. derived and harvested organoids for mass spectrometry, imaging, and qPCR assays. A.L. performed and analyzed qPCR data. L.E.A.G. conducted electrophysiological recordings of organoids. S.M. and K.W. performed optical clearing, immunostainings, and imaging of organoids with help from A.L.. S.M. designed and optimized multiome experiments with inputs from S. and C.M.S.. S.M. and S. processed the organoids for multiomic sequencing. A.C. and S.E.B. provided pre-processed genetic data. S.M. analyzed multiome sequencing and proteomic data. S.M., S., and C.M.S. interpreted the data. S.M., S., and C.M.S. wrote and edited the manuscript with inputs from co-authors.

