## Extended Data Figures for "A schizophrenia-associated radial glia cell state perturbs early human brain development"

**Extended Data Figure 1**

**(A)** Schematic representation of microglia-containing forebrain organoid generation protocol[18]. Days are indicated as 'D'. EBs: Embryoid bodies, Pre-EMP: Pre-eythromyeloid progenitors, YS: Yolk sac, PMP:pre-macrophage precursors, NPC: Neural precursor cells, iPSCs: induced pluripotent stem cells. **(B)** Brightfield microscope images (4X magnification) of individual steps during microglia-containing forebrain organoid generation starting from patient-derived iPSCs to maturing organoids at DIV 120 (left to right). **(C)** Confocal microscopy images of immunohistochemical staining for neurodevelopmental markers on organoid sections (18μm). **(D)** Electrophysiological membrane properties in live patient-derived organoids by whole-cell patch clamping. Representative trace of firing response to a square voltage step from an organoid per condition (top; unaffected $MZ_d$ in cyan, Affected $MZ_d$ in magenta). Bar graph of Membrane capacitance (Cm), Membrane potential (mV), Firing threshold, and number of maximum number of action potentials per cell. Data is presented as means + s.e.m. in bar graphs. * Indicates $p<0.05$, **$p<0.01$, ***$p<0.005$. **(E)** Protein-protein interaction network (string-db) of all significantly dysregulated proteins in DIV 120 organoids derived from SCZ-affected twin subjects compared to their respective unaffected twins. Coloured nodes indicates log-fold change values of downregulated proteins (white to green) and upregulated proteins (white to red).

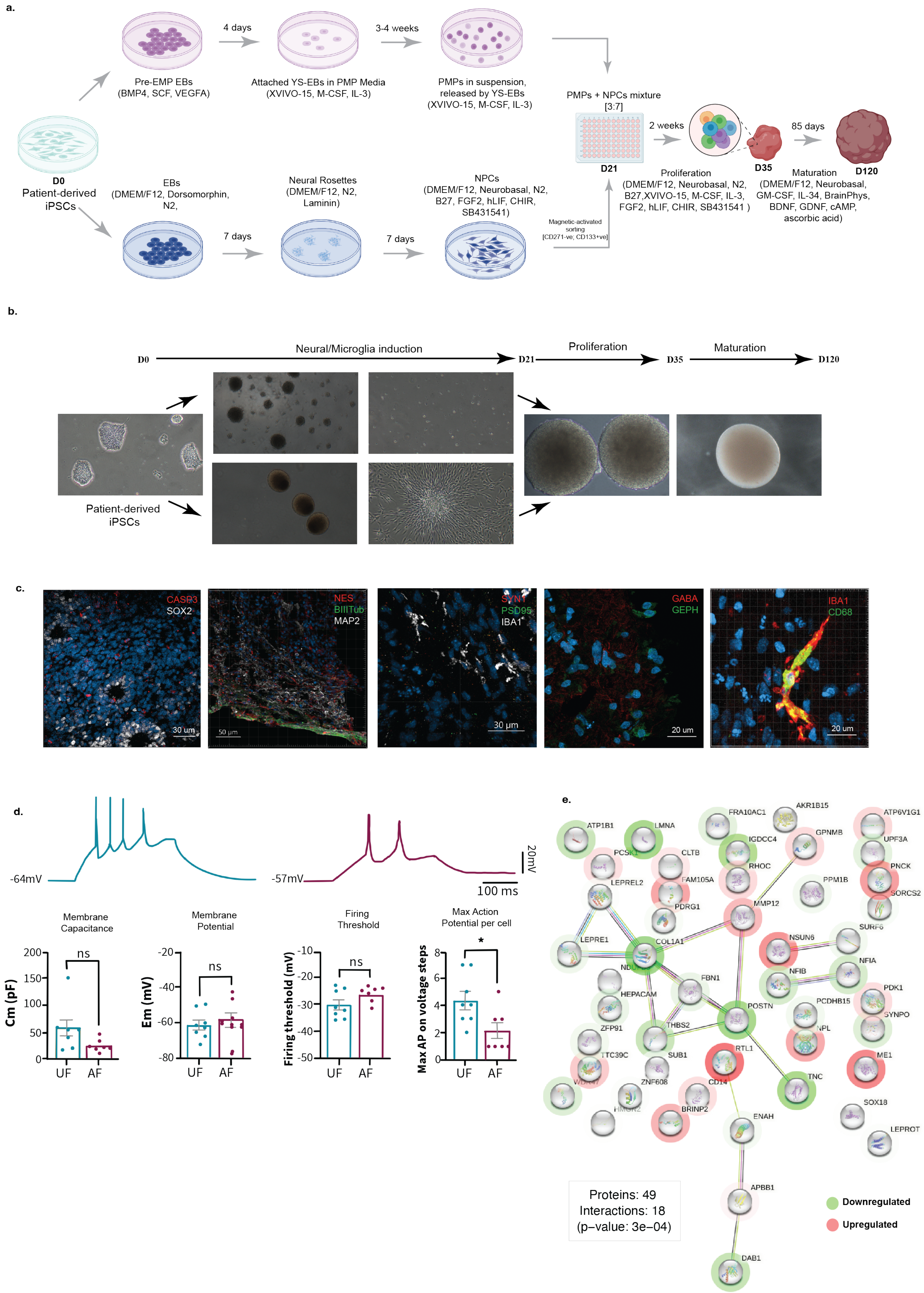
a.
4 days
3-4 weeks
Pre-EMP EBs
(BMP4, SCF, VEGFA)
Attached YS-EBs in PMP Media
(XVIVO-15, M-CSF, IL-3)
PMPs in suspension,
released by YS-EBs
(XVIVO-15, M-CSF, IL-3)
PMPs + NPCs mixture
[3:7]
2 weeks
85 days
D21
D35
D120
D0
Patient-derived
iPSCs
EBs
(DMEM/F12, Dorsomorphin,
N2,
Neural Rosettes
(DMEM/F12, N2,
Laminin)
NPCs
(DMEM/F12, Neurobasal, N2,
B27, FGF2, hLIF, CHIR,
SB431541)
Magnetic-activated
sorting
[CD271-ve; CD133+ve]
7 days
7 days
Proliferation
(DMEM/F12, Neurobasal, N2,
B27,XVIVO-15, M-CSF, IL-3,
FGF2, hLIF, CHIR, SB431541 )
Maturation
(DMEM/F12, Neurobasal,
GM-CSF, IL-34, BrainPhys,
BDNF, GDNF, cAMP,
ascorbic acid)
b.
D0
Neural/Microglia induction
D21
Proliferation
D35
Maturation
D120
Patient-derived
iPSCs
c.
CASP3
SOX2
30 um
NES
BIIITub
MAP2
50 µm
SYN1
PSD95
IBA1
30 µm
GABA
GEPH
20 um
IBA1
CD68
20 um
d.
-64mV
-57mV
20mV
100 ms
Membrane
Capacitance
Membrane
Potential
Firing
Threshold
Max Action
Potential per cell
Cm (pF)
Em (mV)
Firing threshold (mV)
Max AP on voltage steps
ns
ns
ns
*
UF
AF
e.
ATP1B1
LMNA
FRA10AC1
AKR1B15
ATP6V1G1
IGDCC4
GPNMB
UPF3A
PCSK1
CLTB
RHOC
PNCK
LEPREL2
FAM105A
PPM1B
SORCS2
PDRG1
MMP12
SURF6
LEPRE1
COL1A1
NSUN6
NFIA
FBN1
NFIB
HEPACAM
PDK1
POSTN
PCDHB15
SYNPO
ZFP91
THBS2
NPL
TTC39C
SUB1
RTL1
ME1
ZNF608
CD14
TNC
SOX18
BRINP2
ENAH
LEPROT
APBB1
DAB1
Proteins: 49
Interactions: 18
(p–value: 3e–04)
Downregulated
Upregulated

**Extended Data Figure 2.**

**(A)** Quality of nuclei (40x) isolated from dissociated patient-derived organoids before loading the 10X Multiome chip J, showing no nuclei clumping or blebbing. **(B)** Number of cells per sample across both conditions. **(C)** Number of genes versus UMI's captured per cell across samples. **(D)** Violin plots showing distribution of quality control metrics per cell across samples. TSS: Transcription start site **(E)** UMAPs of batch-integrated space across RNA and ATAC data using two computational algorithms, covariate Disentangling Augmented Loss (CODAL) implemented in Mira[64] (left) and weighted nearest-neighbors (WNN) implemented in Seurat[62] (right). Cells are colored by samples. **(F)** UMAP distribution of metadata of subject cells (twin pair identifier, condition, sex and sample), and QC metrics (percentage of mitochondrial transcripts, G2M cell cycle phase score, percentage of ribosomal transcripts, RNA transcripts, RNA genes, ATAC fragments, ATAC-seq peaks and ratio of mononucleosomal to nucleosome-free fragments). **(G)** UMAP of computed clustering using the Louvain algorithm on codal-integrated joint-KNN graph. **(H)** Composition of annotated clusters across subjects. **(I)** Heatmap of Pearson's correlations of gene expression between computed clusters. **(J)** Fragment length periodicity of all cells with percentage of fragments (y axis) and fragment size in base pairs (x-axis). **(K)** Density plot showing TSS enrichment score (y-axis) versus unique fragments per cell (x-axis) prior to filtering. Dotted line indicates TSS enrichment cutoff.

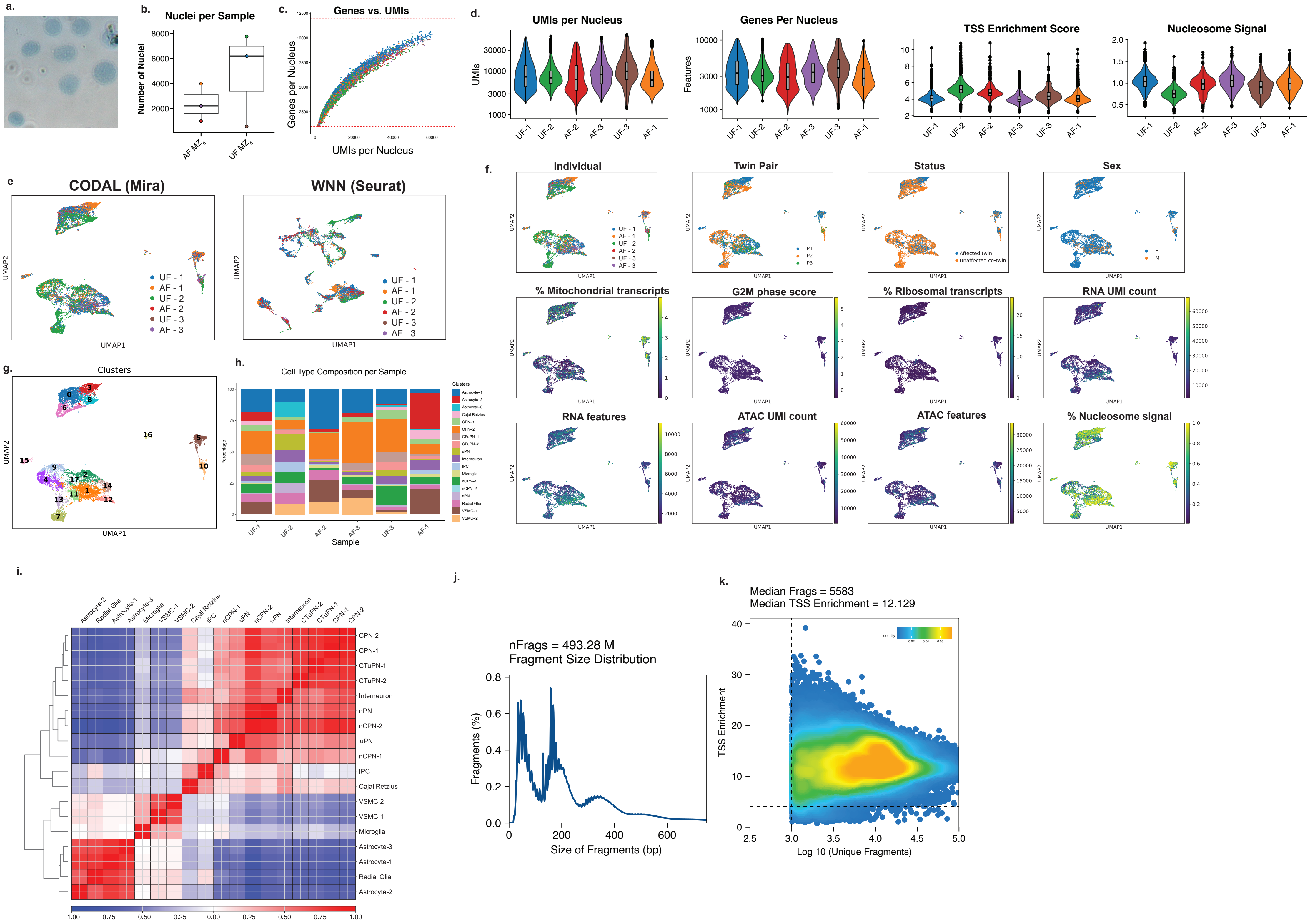

a.
b.
Nuclei per Sample
Number of Nuclei
AF MZ
UF MZ
c.
Genes vs. UMIs
Genes per Nucleus
UMIs per Nucleus
d.
UMIs per Nucleus
UMIs
Genes Per Nucleus
Features
TSS Enrichment Score
Nucleosome Signal
UF-1
UF-2
AF-2
AF-3
UF-3
AF-1
e
CODAL (Mira)
WNN (Seurat)
UMAP1
UMAP2
UF - 1
AF - 1
UF - 2
AF - 2
UF - 3
AF - 3
f.
Individual
Twin Pair
P1
P2
P3
Status
Affected twin
Unaffected co-twin
Sex
F
M
% Mitochondrial transcripts
G2M phase score
% Ribosomal transcripts
RNA UMI count
RNA features
ATAC UMI count
ATAC features
% Nucleosome signal
g.
Clusters
h.
Cell Type Composition per Sample
Percentage
Sample
Clusters
Astrocyte-1
Astrocyte-2
Astrocyte-3
Cajal Retzius
CPN-1
CPN-2
CFuPN-1
CFuPN-2
Interneuron
uPN
IPC
Microglia
nCPN-1
nCPN-2
nPN
Radial Glia
VSMC-1
VSMC-2
i.
Astrocyte-2
Radial Glia
Astrocyte-1
Astrocyte-3
Microglia
VSMC-1
VSMC-2
Cajal Retzius
IPC
nCPN-1
uPN
nCPN-2
nPN
Interneuron
CTuPN-2
CTuPN-1
CPN-1
CPN-2
j.
nFrags = 493.28 M
Fragment Size Distribution
Fragments (%)
Size of Fragments (bp)
k.
Median Frags = 5583
Median TSS Enrichment = 12.129
density
TSS Enrichment
Log 10 (Unique Fragments)

**Extended Data Figure 3.**

UMAP plots depicting the distribution of all identified **(A)** expression and **(B)** accessibility topic activation scores per cell.

a.

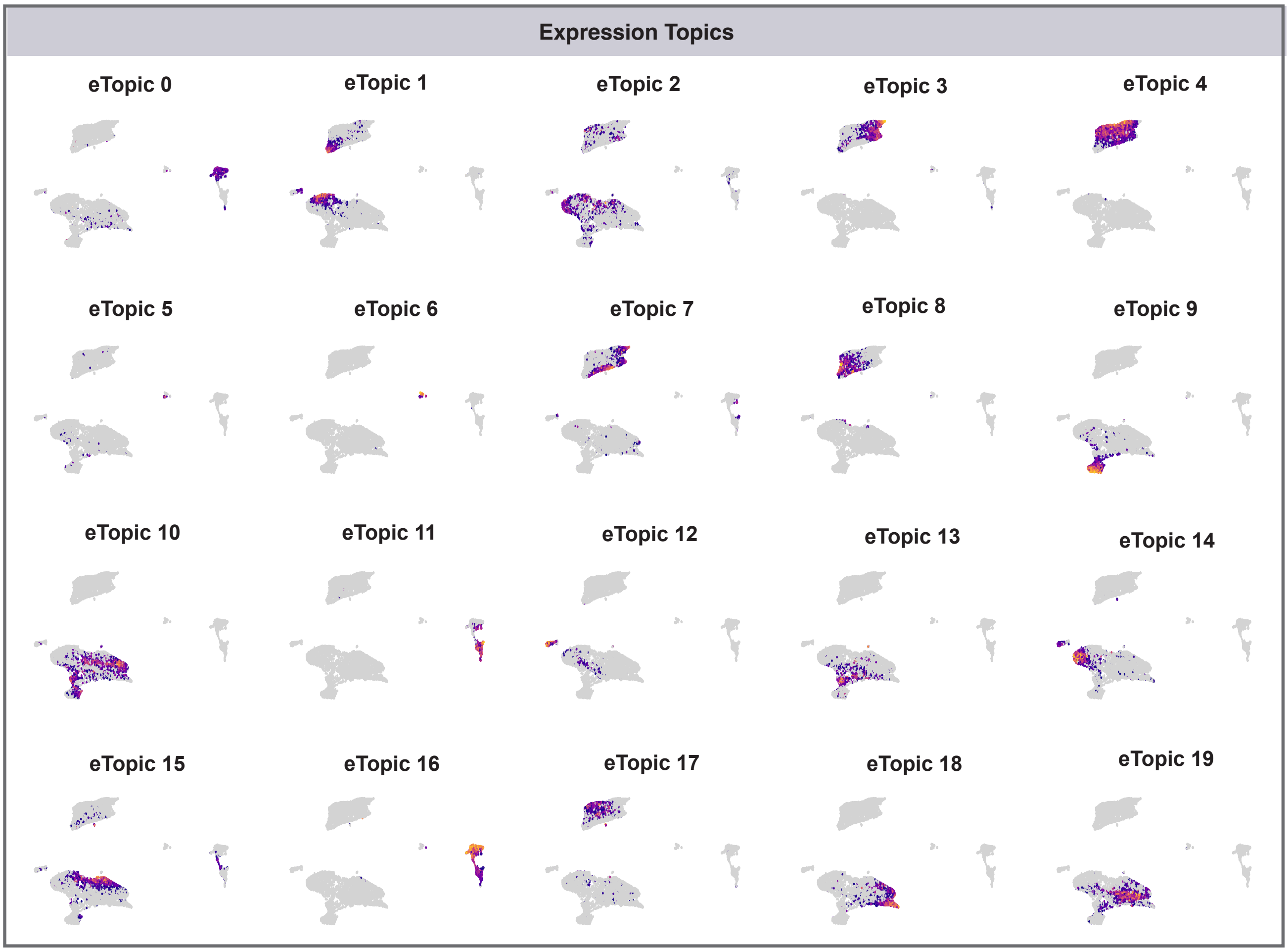
Expression Topics
eTopic 0
eTopic 1
eTopic 2
eTopic 3
eTopic 4
eTopic 5
eTopic 6
eTopic 7
eTopic 8
eTopic 9
eTopic 10
eTopic 11
eTopic 12
eTopic 13
eTopic 14
eTopic 15
eTopic 16
eTopic 17
eTopic 18
eTopic 19

b.

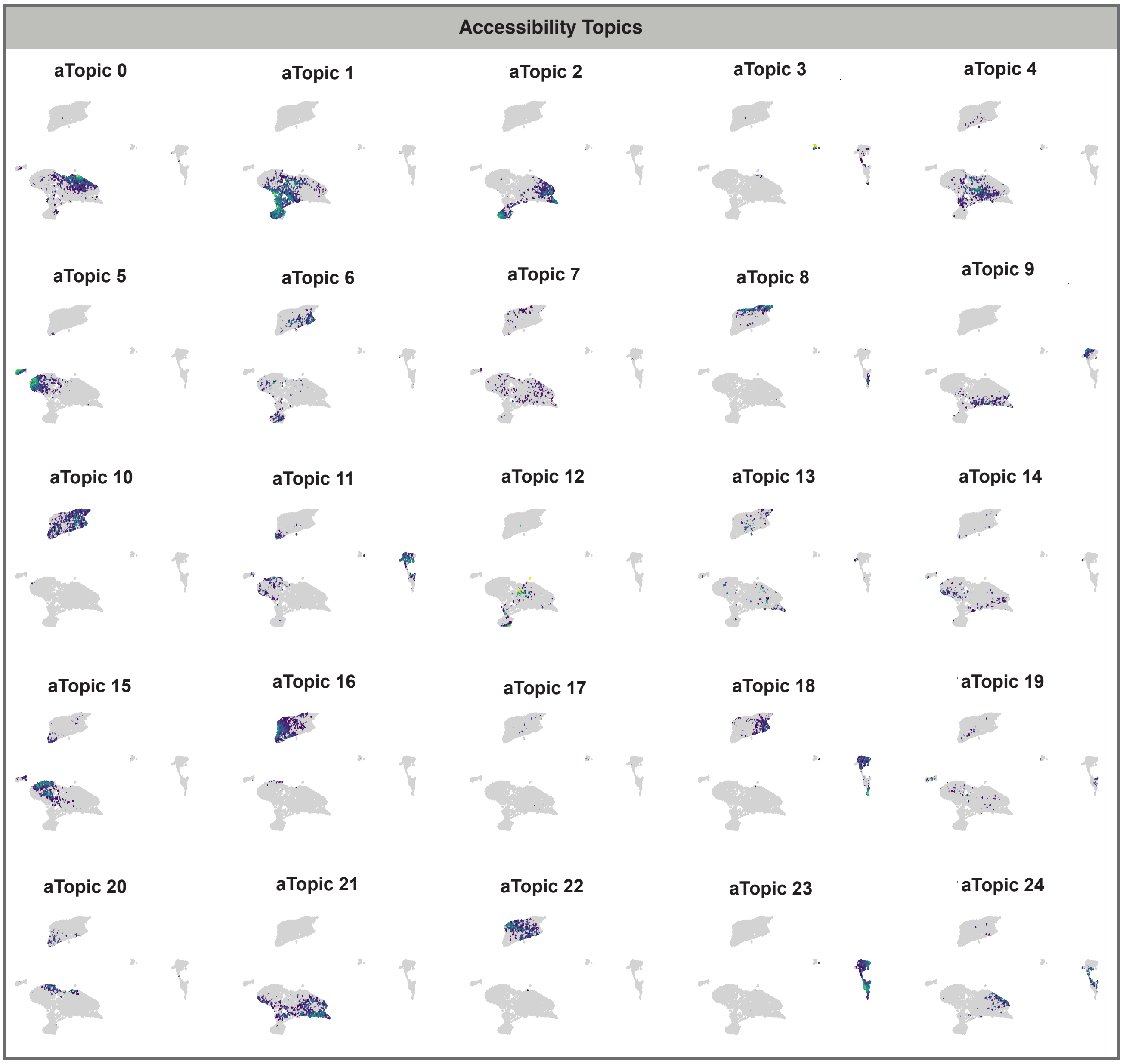
Accessibility Topics
aTopic 0
aTopic 1
aTopic 2
aTopic 3
aTopic 4
aTopic 5
aTopic 6
aTopic 7
aTopic 8
aTopic 9
aTopic 10
aTopic 11
aTopic 12
aTopic 13
aTopic 14
aTopic 15
aTopic 16
aTopic 17
aTopic 18
aTopic 19
aTopic 20
aTopic 21
aTopic 22
aTopic 23
aTopic 24

**Extended Data Figure 4.**

**(A)** Multimodal reference mapping using unsupervised label transfer to map organoid clusters to multimodal data (expression and accessibility measured within the same cell) from primary fetal brain during development and postnatal lifespan, generated using the same platform[76]. UMAPs display cell type prediction scores assigned to each cell. Blue color scale indicates age predictions while red scale indicates cell type predictions. Prediction scores are computed between 0 to 1. **(B)** Cluster correlations between organoid clusters (RNA modality) and fetal brain atlases generated from human fetal brain tissue at multiple periods[77–80]. Color scale denotes pearson's correlation coefficient. **(C)** Cluster correlations of individual organoid clusters (top) and whole organoid (bottom) (RNA modality) to BrainSpan Atlas (https://www.brainspan.org/) from fetal brain tissue spanning the entire lifespan (bulk RNA-seq). Color scale denotes pearson's correlation coefficient. **(D)** Dot plot visualizing the distribution of cluster-specific gene expression markers from the RNA modality. Color scale denotes normalized expression levels and size indicated the percentage of cells expressing the marker. **(E)** Heatmap of computed TF-binding motif (y-axis) scores based on identified cluster-specific accessible chromatin regions across cells grouped by cell types (x-axis).

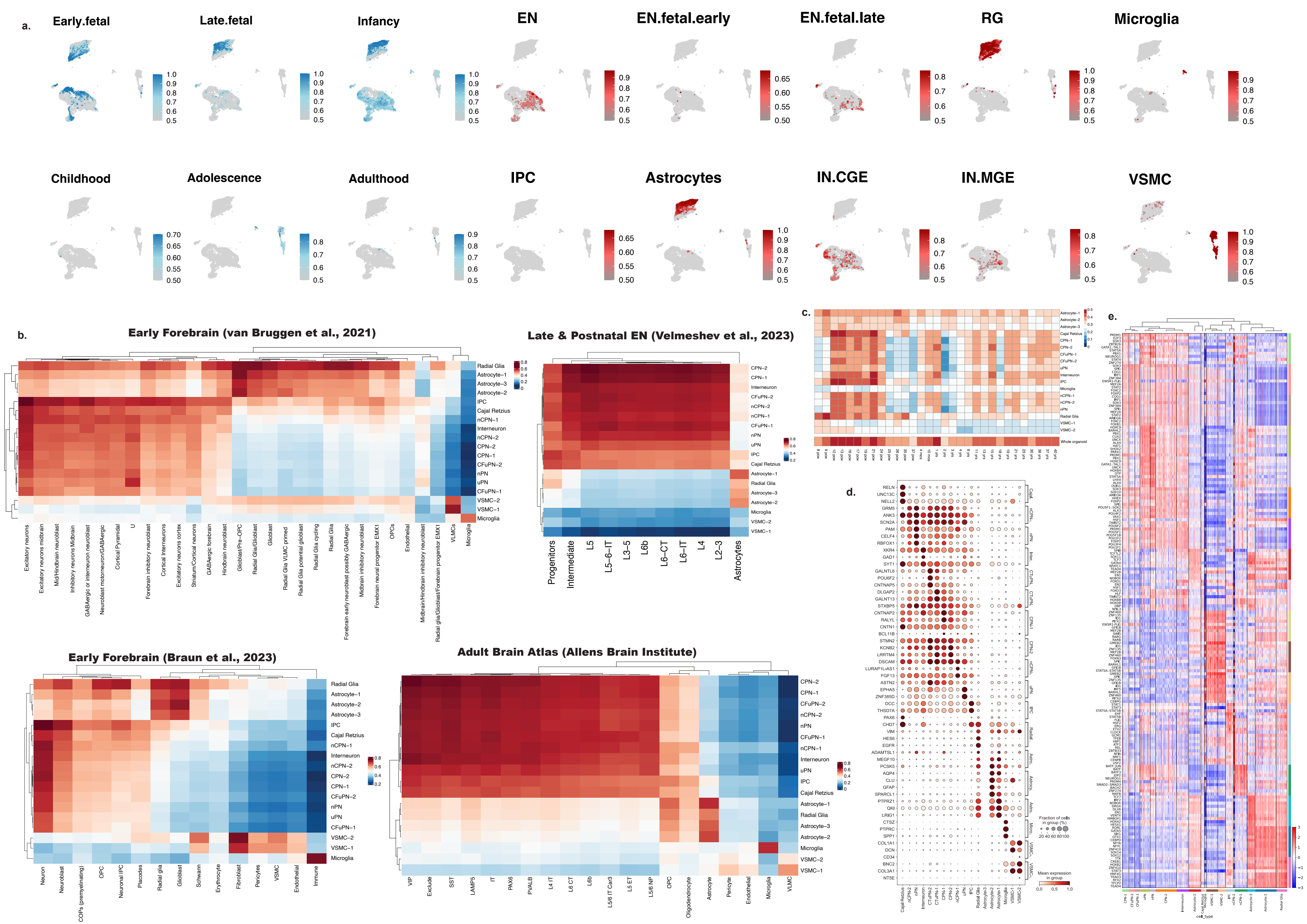
a.
Early.fetal
Late.fetal
Infancy
EN
EN.fetal.early
EN.fetal.late
RG
Microglia
Childhood
Adolescence
Adulthood
IPC
Astrocytes
IN.CGE
IN.MGE
VSMC
b.
Early Forebrain (van Bruggen et al., 2021)
Late & Postnatal EN (Velmeshev et al., 2023)
Early Forebrain (Braun et al., 2023)
Adult Brain Atlas (Allens Brain Institute)
Radial Glia
Astrocyte−1
Astrocyte−2
Astrocyte−3
IPC
Cajal Retzius
nCPN−1
nCPN−2
Interneuron
CPN−1
CPN−2
CFuPN−1
CFuPN−2
nPN
uPN
VSMC−1
VSMC−2
Microglia
Progenitors
Intermediate
L5
L5−6−IT
L3−5
L6b
L6−CT
L6−IT
L4
L2−3
Astrocytes
Neuron
Neuroblast
COPs (premyelinating)
OPC
Neuronal IPC
Placodes
Radial glia
Glioblast
Schwann
Erythrocyte
Fibroblast
Pericytes
VSMC
Endothelial
Immune
VIP
Exclude
SST
LAMP5
IT
PAX6
PVALB
L4 IT
L6 CT
L6b
L5/6 IT Car3
L5 ET
L5/6 NP
OPC
Oligodendrocyte
Astrocyte
Pericyte
Endothelial
Microglia
VLMC
c.
Whole organoid
d.
Fraction of cells in group (%)
20 40 60 80 100
Mean expression in group
0.0 0.5 1.0
e.
cell_type

**Extended Data Figure 5**

**(A)** Branch-specific genes tested for association with each trajectory using generalized additive models (GAM) models (P-adjusted<0.05). Fitted genes are smoothed over pseudotime, clustered into modules and visualized combined on UMAPs (top) coloured by computed fate probabilities per cell and individually on heatmaps (bottom) across pseudotime (x-axis) per branch leading to a terminal fate. **(B)** UMAP embedding of the recontructed joint KNN graph with cells colored by identified cell types. **(C)** Compositional data analysis loadings (x-axis) for cell types (y-axis) within the organoids compared across conditions. Positive loadings correspond to over-representation in the affected $MZ_d$ and negative loadings mean over-representation in unaffected $MZ_d$. Boxplots (left) depict the uncertainty of the loading coefficients under cell and individual subsampling. Bar plots on the left depict log-transformed p-values (x-axis). Black line depicts Bonferroni threshold 0.01. **(D)** UMAP embedding with cells colored according to their sample-associated relative SCZ-likelihood scores calculated using the MELD algorithm. **(E)** Significant ligand-receptor interactions, inferred between astrocytes, radial glia and neuronal clusters (x-axis), that are significantly upregulated in SCZ-affected $MZ_d$ compared to unaffected $MZ_d$. Color scale represents inferred communication probability score. Size of the dot represents significance of inferred communication between the ligand-receptor pair (y-axis). **(F)** Pie chart displaying the proportion of genetic elements across discordant regions identified as deleted (left) or duplicated (right) (<=4) within nine twin pairs (n=18 subjects) from WGS data of the independent cohort. **(G)** Enrichment of biological pathways (y-axis; GO:BP) within the overlapping transcriptional networks of *SORCS3* and *SORCS2* within the developing brain. Individual seeded-transcriptional networks for *SORCS2* and *SORCS3* were constructed using publicly available RNA-seq data of the developing brain (BrainSpan database). Postive networks include genes positively correlated to the respective receptor whereas negative networks include genes negatively correlated to the receptor. The enrichment was tested on genes overlapping between resepctive networks of the two receptors. p-values were adjusted for multiple testing using Benjamini Hochberg-method. Number of genes in each pathway contributing to the enrichement is indicated in white.

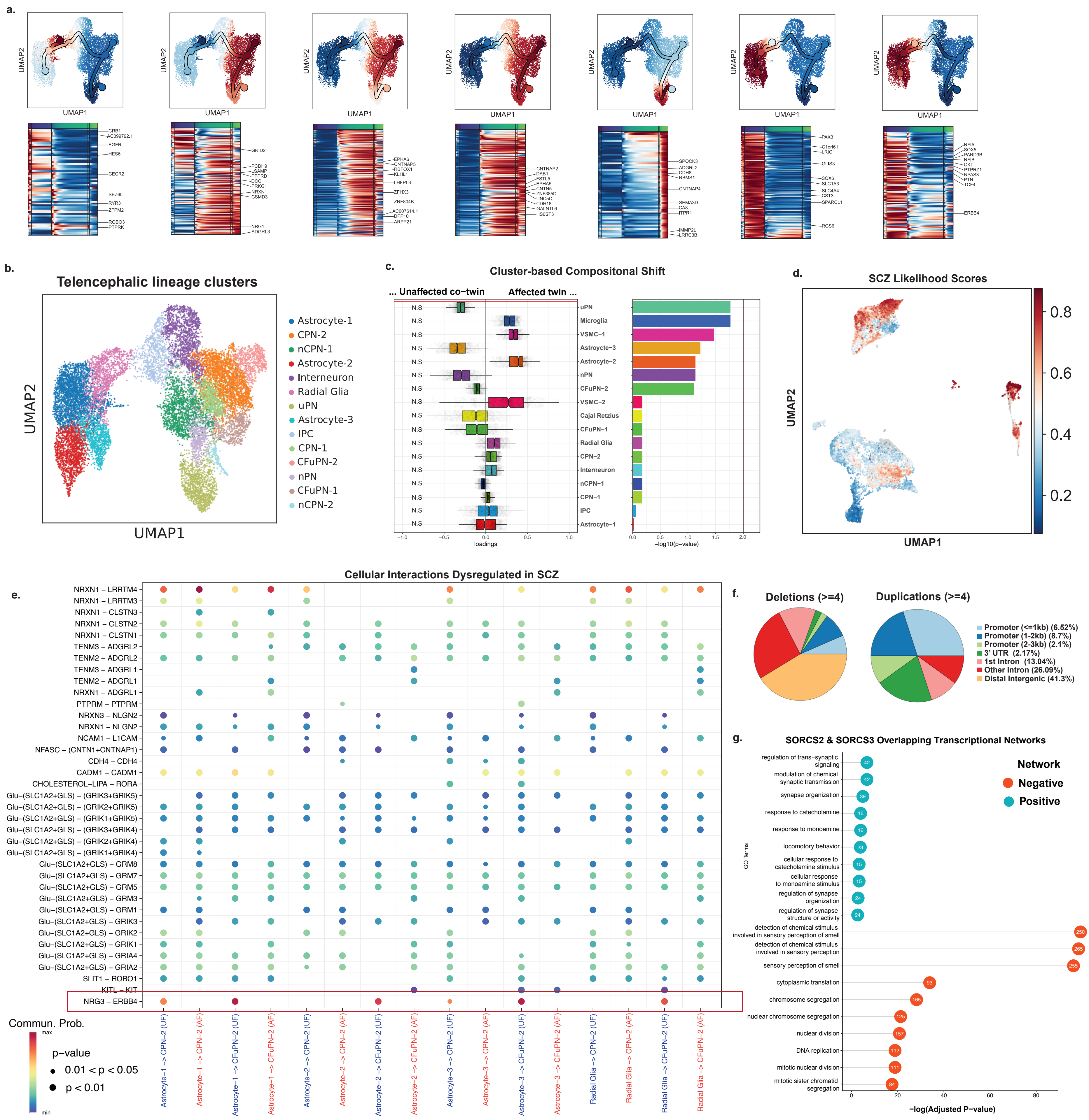
a.
UMAP1
UMAP2
b.
Telencephalic lineage clusters
Astrocyte-1
CPN-2
nCPN-1
Astrocyte-2
Interneuron
Radial Glia
uPN
Astrocyte-3
IPC
CPN-1
CFuPN-2
nPN
CFuPN-1
nCPN-2
c.
Cluster-based Compositonal Shift
... Unaffected co−twin
Affected twin ...
loadings
−log10(p-value)
d.
SCZ Likelihood Scores
e.
Cellular Interactions Dysregulated in SCZ
Receptor-Ligand Pair
Commun. Prob.
p−value
0.01 < p < 0.05
p < 0.01
f.
Deletions (>=4)
Duplications (>=4)
Promoter (<=1kb) (6.52%)
Promoter (1-2kb) (8.7%)
Promoter (2-3kb) (2.1%)
3' UTR (2.17%)
1st Intron (13.04%)
Other Intron (26.09%)
Distal Intergenic (41.3%)
g.
SORCS2 & SORCS3 Overlapping Transcriptional Networks
Network
Negative
Positive
GO Terms
−log(Adjusted P−value)

**Extended Data Figure 6.**

**(A)** Density distribution of cells belonging to each condition along the progression of glial lineage as indiciated by pseudotime (y-axis). **(B)** Heatmap showing mean smoothed expression values of significant trajectory-associated differentially expressed genes (pseudotime*condition) in the glial lineage across pseudotime (y-axis) (P-adjusted < 0.05). Genes are clustered using heirarchical clustering. Red box highlights genes involved in cell cycle proliferation dysregulated in SCZ-affected twins. **(C)** Scatter plot of log-transformed normalized expression (y-axis) of selected trajectory-associated DEGs across pseudotime (x-axis) in the glial lineage. Cells are coloured according to the condition as shown in the legend. Colored line represents a smoothing spline fit in the respective condition. **(D)** PHATE embedding of Astrocyte-1 (AST-1) cluster with cells colored by the identified vertex frequency subclusters and **(E)** SCZ-likelihood scores. **(E)** Jitter plot displays cells belonging to each $MZ_d$ subject across both AST-1 cell states (x-axis) and their SCZ-likelihood scores (y-axis). AST-1 cell state enriched in SCZ-affected $MZ_d$ is labelled as scz-AST-1 and unaffected $MZ_d$ is labelled as $_n$scz-AST-1. State that exhibits cells from both conditions is labelled as mAST-1. Legend indicates the $MZ_d$ twin subject. Top significant pathways of SCZ-enriched **(G)** scz-AST-2 and **(H)** scz-AST-1 cell states.

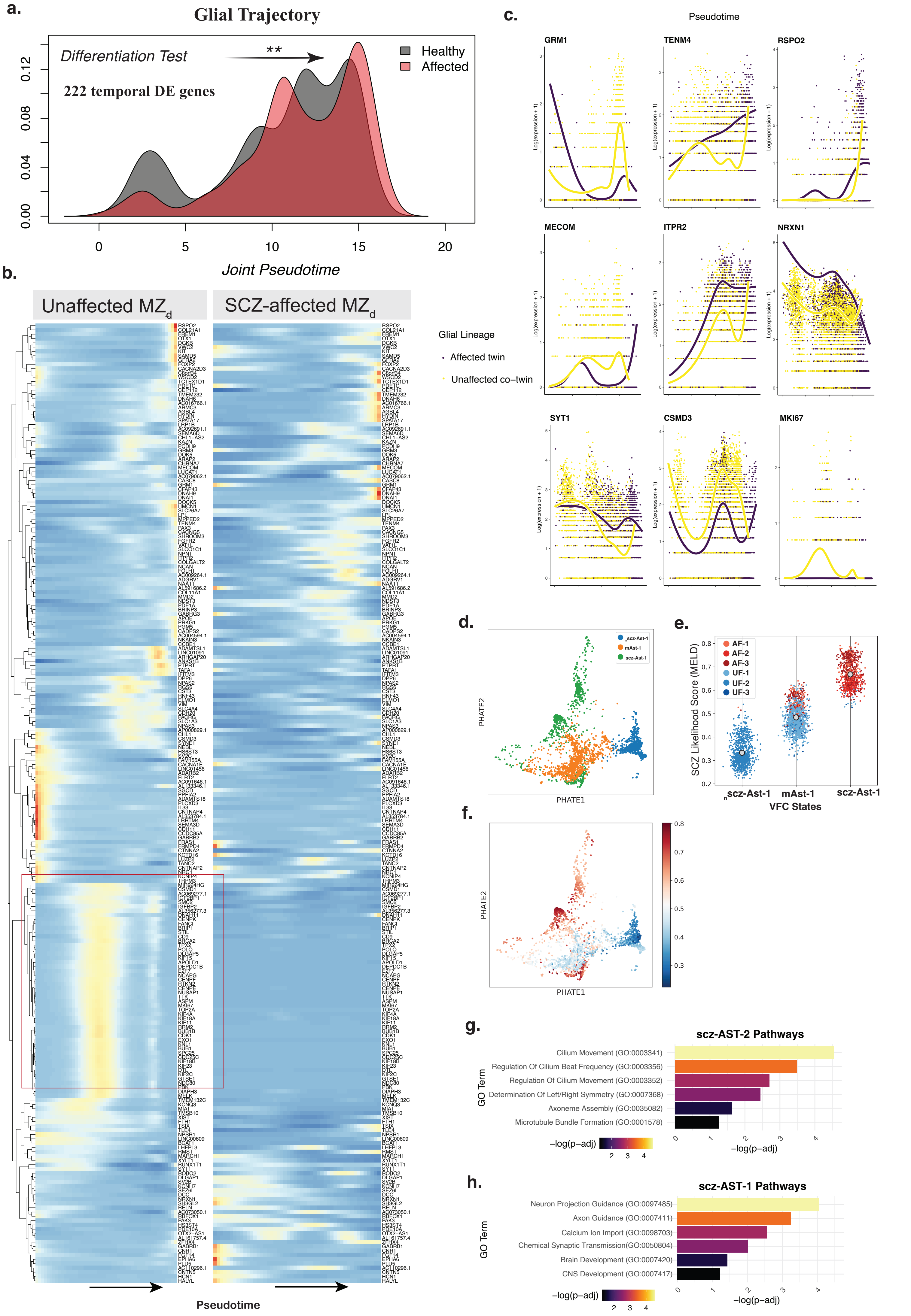
a.
Glial Trajectory
Differentiation Test
**
222 temporal DE genes
Healthy
Affected
0.00
0.04
0.08
0.12
0
5
10
15
20
Joint Pseudotime
b.
Unaffected MZd
SCZ-affected MZd
Pseudotime
c.
Pseudotime
GRM1
TENM4
RSPO2
MECOM
ITPR2
NRXN1
SYT1
CSMD3
MKI67
Log(expression + 1)
Glial Lineage
Affected twin
Unaffected co−twin
d.
PHATE2
PHATE1
scz-Ast-1
mAst-1
scz-Ast-1
e.
AF-1
AF-2
AF-3
UF-1
UF-2
UF-3
SCZ Likelihood Score (MELD)
scz-Ast-1
mAst-1
scz-Ast-1
VFC States
f.
PHATE2
PHATE1
g.
scz-AST-2 Pathways
Cilium Movement (GO:0003341)
Regulation Of Cilium Beat Frequency (GO:0003356)
Regulation Of Cilium Movement (GO:0003352)
Determination Of Left/Right Symmetry (GO:0007368)
Axoneme Assembly (GO:0035082)
Microtubule Bundle Formation (GO:0001578)
GO Term
−log(p−adj)
h.
scz-AST-1 Pathways
Neuron Projection Guidance (GO:0097485)
Axon Guidance (GO:0007411)
Calcium Ion Import (GO:0098703)
Chemical Synaptic Transmission(GO:0050804)
Brain Development (GO:0007420)
CNS Development (GO:0007417)
GO Term
−log(p−adj)
